# Statistically Consistent Rooting of Species Trees under the Multispecies Coalescent Model

**DOI:** 10.1101/2022.10.26.513897

**Authors:** Yasamin Tabatabaee, Sebastien Roch, Tandy Warnow

## Abstract

Rooted species trees are used in several downstream applications of phylogenetics. Most species tree estimation methods produce unrooted trees and additional methods are then used to root these unrooted trees. Recently, Quintet Rooting (QR) (Tabatabaee et al., ISMB and Bioinformatics 2022), a polynomial-time method for rooting an unrooted species tree given unrooted gene trees under the multispecies coalescent, was introduced. QR, which is based on a proof of identifiability of rooted 5-taxon trees in the presence of incomplete lineage sorting, was shown to have good accuracy, improving over other methods for rooting species trees when incomplete lineage sorting was the only cause of gene tree discordance, except when gene tree estimation error was very high. However, the statistical consistency of QR was left as an open question. Here, we present QR-STAR, a polynomial-time variant of QR that has an additional step for determining the rooted shape of each quintet tree. We prove that QR-STAR is statistically consistent under the multispecies coalescent model. Our simulation study under a variety of model conditions shows that QR-STAR matches or improves on the accuracy of QR. QR-STAR is available in open source form at https://github.com/ytabatabaee/Quintet-Rooting.

## 1 Introduction

Inferring rooted species trees is important for many downstream applications of phylogenetics, such as comparative genomics [11,7] and dating [26]. These estimations use different loci from across the genomes of the selected species, and so are referred to as multi-locus analyses. If rooted gene trees can be accurately inferred, then the rooted species tree can be estimated from them [14]; however, this is not a reliable assumption [30]. Hence, the standard approach is to first estimate an unrooted species tree using multi-locus datasets, and then root that estimated tree.

The estimation of the unrooted species tree is challenged by biological processes, such as incomplete lineage sorting (ILS) or gene duplication and loss (GDL), that can result in different parts of the genome having different evolutionary trees. When ILS or GDL occur, statistically consistent estimation of the unrooted species tree requires techniques that take the source of heterogeneity into consideration [15,23]. The case of ILS, as modeled by the multispecies coalescent (MSC) model [10], is the most well-studied, and there are several methods for estimating unrooted species trees that have been proven statistically consistent under the MSC (see [23] for a survey of such methods).

The general problem of rooting a species tree (or indeed even a gene tree) is of independent interest, but presents many challenges. A common approach is the use of an outgroup taxon (i.e., the inclusion of a species that is outside the smallest clade containing the remaining species), so that the resultant tree is rooted on the edge leading to the outgroup [16]. However, outgroup selection has its own difficulties: if the outgroup is too distant, then it may be attached fairly randomly to the tree containing the remaining species, and if it is too close, it may even be an ingroup taxon [5,6,9,13]. Other approaches use branch lengths estimated on the tree to find the root based on specific optimization criteria; however, these approaches tend to degrade in accuracy unless the strict molecular clock holds (which assumes that all sites along the genome evolve under a constant rate) [8,18,32].

Quintet Rooting (QR) [31] is a recently introduced method that is designed to root a given species tree using the unrooted gene tree topologies, under the assumption that the gene trees can differ from the species tree due to ILS. QR is based on mathematical theory established by Allman, Degnan, and Rhodes [2], which showed that the rooted species tree topology is identifiable from the unrooted gene tree topologies whenever the number of species is at least five. In [31], QR was shown to provide good accuracy for rooting both estimated and true species trees in the presence of ILS, compared to alternative methods.

However, QR was not proven to be statistically consistent–which addresses the theoretical question of whether the root location selected by QR, when given the true species tree, will converge to the correct location as the number of gene trees in the input increases. Although much attention has been paid to establishing statistical consistency for unrooted species tree estimation methods and many methods, such as ASTRAL [19], SVDQuartets [33] and BUCKy [12] have been proven to be statistically consistent estimators of the unrooted species tree topology under the MSC, to the best of our knowledge, no prior study has addressed the statistical consistency properties of methods for rooting species trees.

In this paper, we argue that QR is not guaranteed to be statistically consistent under the MSC, but we also present a modification to QR, which we call QR-STAR, that we prove statistically consistent. Moreover, QR-STAR, like QR, runs in polynomial time.

The rest of this paper is organized as follows: Section 2 provides background information on QR, including the theory established by Allman, Degnan, and Rhodes [2]. Section 3 introduces QR-STAR and Section 4 provides the proof of statistical consistency of QR-STAR under the MSC. In Section 5 we present the results of a simulation study exploring the parameter space of QR-STAR and evaluating its performance in comparison to QR. We conclude in Section 6. Due to space limitations, most of the proofs and results from the simulation study are presented in the Supplementary Materials.

## 2 Background

We present the theory from [2] first, which establishes identifiability of the rooted species tree from unrooted quintet trees, and then we describe Quintet Rooting (QR), our earlier method for rooting species trees. Together these form the basis for deriving our new method, QR-STAR, which we present in the next section.

### 2.1 Allman, Degnan, and Rhodes (ADR) Theory

Allman, Degnan, and Rhodes (ADR) [2] established that the unrooted topology of the species tree is identifiable from four-leaf unrooted gene trees under the MSC, a result that is well known and used in several “quartet-based” methods for estimating species trees under the MSC [12,19,17,25]. ADR also proved that the rooted species tree topology is identifiable from unrooted five-leaf gene tree topologies; this result is much less well known, but was recently used in the development of QR for rooting species trees.

ADR have described the probability distribution of unrooted gene tree topologies under each 5-taxon MSC model species tree. On a given set of five taxa, there exist 105 different rooted binary trees, labeled with *R*_1_, …, *R*_105_^3^, that can be categorized into three groups based on their (unlabeled) rooted shapes: caterpillar, balanced and pseudo-caterpillar [28]. An example of a tree from each category is shown in Figure 1. Each 5-taxon model species tree defines a specific probability distribution over the 15 different unrooted gene tree topologies on the same leafset, shown with *T*_1_, …, *T*_15_. Theorem 9 in [2] states that this distribution uniquely determines the rooted tree topology and its internal branch lengths for trees with at least five taxa.

To prove this identifiability result, the ADR theory specifies a set of linear invariants (i.e., equalities) and inequalities that must hold between the probabilities of unrooted 5-taxon gene trees, for any choice of the parameters of the model species tree. These linear invariants and inequalities define a partial order on the distribution of 5-taxon unrooted gene tree topologies. In other words, two gene tree probabilities *u*_*i*_ = ℙ(*T*_*i*_) and *u*_*j*_ = ℙ(*T*_*j*_) can have one of four possible relationships: *u*_*i*_ *> u*_*j*_, *u*_*j*_ *> u*_*i*_, *u*_*i*_ = *u*_*j*_, or *u*_*i*_ and *u*_*j*_ are not comparable.

**Fig. 1:**
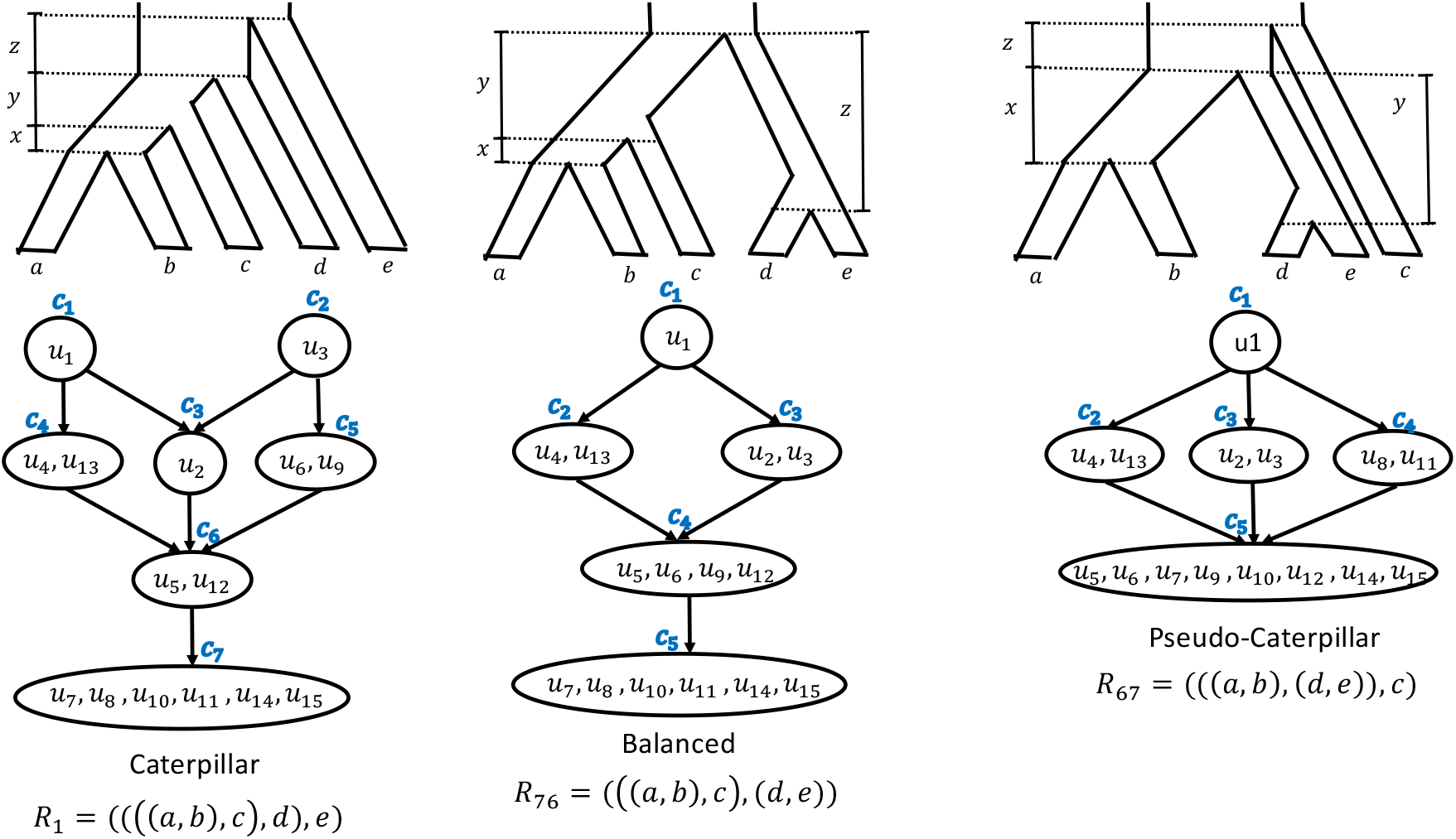
ADR invariants and inequalities for different rooted topological shapes. The invariants and inequalities found by ADR define a partial order on the distribution of unrooted gene trees for rooted 5-taxon model species trees with different rooted shapes (caterpillar, balanced and pseudo-caterpillar). There are 15 unrooted binary trees on a given set of 5 leaves. Each of the 105 5-taxon rooted species trees define a specific distribution on the probabilities of these unrooted trees. The topology of the rooted binary species tree can be determined from this distribution (i.e., it is identifiable, as established by ADR). While the branch lengths of the rooted species tree depend on the actual probabilities, the linear invariants and inequalities that hold for these distributions is enough to determine the rooted topology of the model species tree.

Figure 1 shows examples of these partial orders with a Hasse diagram for a particular leaf labeling of trees from each rooted shape. Note that some probabilities are members of the same set (e.g., for *R*_1_, set *c*_4_ contains both *u*_4_ and *u*_13_, indicating that *u*_4_ = *u*_13_), and so we refer to the sets *c*_*i*_ as equivalence classes on these probabilities. Furthermore, we will denote the set of equivalence classes associated with a 5-taxon rooted tree *R* with *C*_*R*_. As it can be seen in Figure 1, the number of equivalence classes of caterpillar, balanced and pseudo-caterpillar trees is 7, 5 and 5 respectively. Each directed edge between two equivalence classes in these Hasse diagrams defines an inequality, so that all gene tree probabilities in class *c*_*a*_ at the source of an edge are greater than all gene tree probabilities in class *c*_*b*_ at the target, and we show this by *c*_*a*_ *> c*_*b*_. The exact values of the unrooted gene tree probabilities depend on the internal branch lengths of the model tree, and ADR provide a set of formulas that relate the model tree parameters to the probability distribution of the unrooted gene trees in Appendix B of [2], which will be used in our proofs.

### 2.2 Quintet Rooting

The input to QR is an unrooted species tree *T* with *n* leaves and a set 𝒢 of *k* single-copy unrooted gene trees where the gene trees draw their leaves from the leafset of *T*, denoted by ℒ(*T*). Given this input, QR searches over all possible rootings of *T* and returns a tree most consistent with the distribution of quintets (i.e., 5-taxon trees) in the input gene trees.

QR approaches this problem by selecting a set *Q* of quintets of taxa from ℒ(*T*) (called the “quintet sampling” step; refer to Appendix A for details), and scoring all rooted versions of *T* based on their induced trees on these quintets. The subtree *T*_|*q*_, *T* restricted to taxa in quintet set *q*, can be rooted on any of its seven edges. In a preprocessing step, QR computes a score for each of these seven different rootings for all trees induced on the quintets in set *Q*, based on a cost function. This results in 7 × |*Q*| computations, and therefore the preprocessing step takes *O*(*k*(|*Q*| + *n*)). Next, for every rooted version of *T*, QR sums up the costs of all its induced rooted trees on quintets in *Q* using the scores computed in the preprocessing step, and returns the rooting with the minimum overall cost. Since *T* can be rooted on any of its 2*n* − 3 edges, the scoring step takes *O*(*n* + |*Q*|) time. Therefore, the overall runtime of QR when using an *O*(*n*) sampling of quintets is *O*(*nk*). Figure 2 shows the pipeline of QR and its individual steps.

**Fig. 2:**
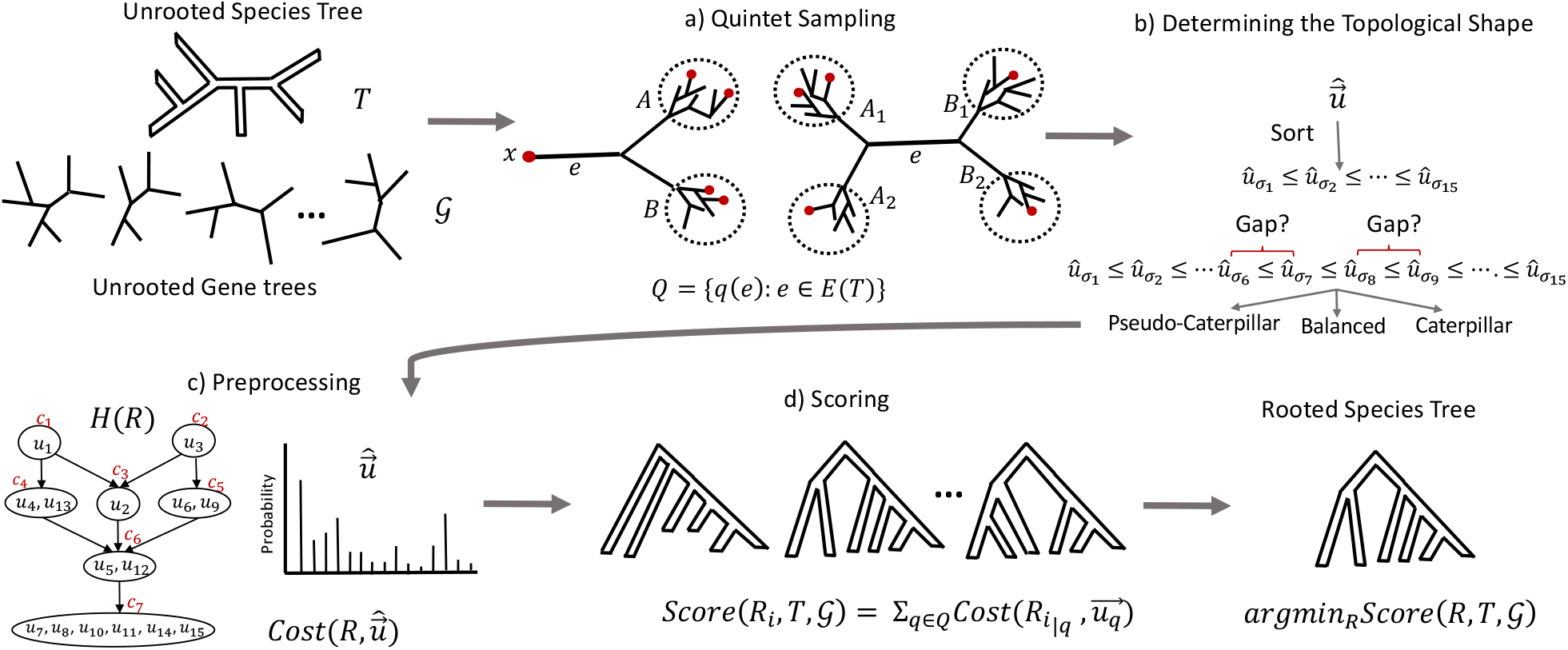
Pipeline of QR and QR-STAR. The input is an unrooted species tree *T* and set of unrooted gene trees 𝒢 on the same leafset. a) The sampling step selects a set *Q* of quintets from the leafset of *T* (shown is the linear encoding sampling). b) An additional step in QR-STAR that determines the rooted shape for each selected quintet. c) The preprocessing step computes a cost for each of the seven possible rootings of each selected quintet. d) The scoring step computes a score for each rooted tree in the search space based on the costs computed in the preprocessing step, and returns a rooting of *T* with minimum score.

Thus, QR provides an exact solution to the optimization problem with the following input and output:

– **Input**: An unrooted tree topology *T*, a set of *k* unrooted gene tree topologies 𝒢 = {*g*_1_, *g*_2_, …, *g*_*k*_}, a set *Q* containing quintets of taxa from leafset ℒ(*T*) and a cost function cost 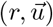.
– **Output**: Rooted tree *R* with topology *T* such that 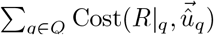 is minimized, where 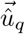 is the distribution of unrooted gene tree quintets in 𝒢|_*q*_ = {*g*_1_|_*q*_, *g*_2_|_*q*_, …, *g*_*k*_|_*q*_}.

#### Cost Function

The cost function 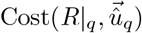 measures the fitness of the rooted quintet tree *R*|_*q*_ with the distribution of the unrooted gene trees restricted to *q* (i.e., 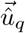), according to the linear invariants and inequalities derived from the ADR theory. In particular, this cost function is designed to penalize a rooted tree *R* _*q*_ if the estimated quintet distribution 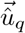 violates some of the inequalities or invariants in its partial order. To this end, a penalty term was considered for each invariant and inequality in the partial order of a 5-taxon rooted tree that is violated in a quintet distribution. The cost function was defined based on a linear combination of these penalty terms, and had the following form, where *r* is a 5-taxon rooted tree and 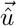 is an estimated quintet distribution:

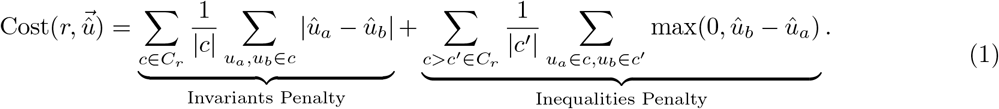

The normalization factors 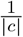 and 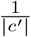 were used to reduce a topological bias that arose from differences in the sizes of the equivalence classes for each tree shape.

## 3 QR-STAR

QR-STAR is an extension to QR that has an additional step for determining the rooted shape (i.e., the rooted topology without the leaf labels) of a quintet tree, as well as an associated penalty term in its cost function. This penalty term compares the rooted shape of the 5-taxon tree, denoted by *S*(*r*), with the rooted shape inferred by QR-STAR from the given quintet distribution, denoted by *Ŝ*(*û*). The motivation for this additional preprocessing step is that, as we argue in Appendix C, the cost function of QR does not guarantee statistical consistency. The cost function of QR-STAR takes the following general form

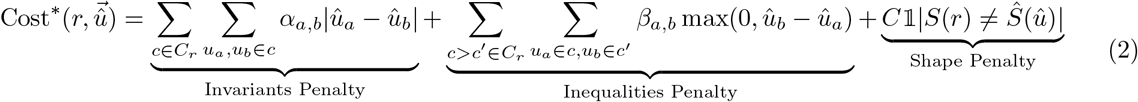

where *α*_*a,b*_ ≥ 0 and *β*_*a,b*_, *C >* 0 are constant real numbers for all *a, b* ^4^. Let *α*_max_ = max_*a,b*_(*α*_*a,b*_) and *β*_min_ = min_*a,b*_(*β*_*a,b*_) where *a, b* ranges over all pairs of indices *a, b* used in the penalty terms in Eq. 2.

Each of the 105 rooted binary trees on a given set of 5 leaves have a unique set of inequalities and invariants that can be derived from the ADR theory. The cost function in Eq. 2 considers a penalty term for these inequalities and invariants as well as the shape of the tree, so that 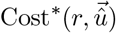 is minimized for a rooted 5-taxon tree *r* that best describes the given estimated quintet distribution.

### 3.1 Determining the Rooted Shape

Model 5-taxon species trees with different rooted shapes (i.e., caterpillar, balanced, pseudo-caterpillar) define equivalence classes with different class sizes on the unrooted gene tree probability distribution. These class sizes can be used to determine the unlabeled shape of a rooted tree, when given the *true* gene tree probability distribution. For example, the size of the equivalence class with the smallest gene tree probabilities is 8 for the pseudo-caterpillar trees and 6 for balanced or caterpillar trees. Therefore, the size of the equivalence class corresponding to the minimal element in the partial order can differentiate a pseudo-caterpillar tree from other tree shapes. Moreover, both caterpillar and balanced trees have a unique class with the second smallest probability, which is of size 2 for caterpillar trees and 4 for balanced trees and this can be used to differentiate a caterpillar tree from a balanced tree. This approach is used in Theorem 9 in [2] for establishing the identifiability of rooted 5-taxon trees from unrooted gene trees.

However, given an *estimated* gene tree distribution, it is likely that none of the invariants derived from the ADR theory exactly hold, and so the class sizes cannot be directly determined and the approach above cannot be used as is to infer the shape of a rooted quintet. Here we propose a simple modification for determining the rooted shape of a tree from the estimated distribution of unrooted gene trees, by looking for significant gaps between quintet gene tree probabilities.

Let *T* be the unrooted species tree with *n* ≥ 5 leaves given to QR-STAR and *q* be a quintet of taxa from) ℒ (*T*). Let 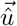 be the quintet distribution estimated from input gene trees induced on taxa in set *q*. QR-STAR first sorts 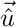 in ascending order to get 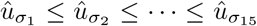. Let 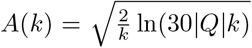 (refer to Lemma 4 for the derivation of *A*(*k*)) where *k* is the number of input gene trees and |*Q*| is the size of the set of sampled quintets in QR-STAR (it depends on the number *n* of taxa and is assumed fixed), and note that lim_*k*→∞_ *A*(*k*) = 0. The first step of QR-STAR computes an estimate of the rooted shape of a quintet *q*, denoted by *Ŝ*(*û*) in Eq. 2, as follows:

– estimate the rooted shape *Ŝ*(*û*) as pseudo-caterpillar if 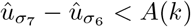;
– estimate the rooted shape *Ŝ*(*û*) as balanced if 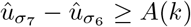 and 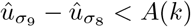;
– estimate the rooted shape *Ŝ*(*û*) as caterpillar if 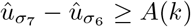 and 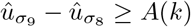.

The runtime of QR-STAR is the same as QR, as determining the topological shape for each quintet is done in constant time, and the overall runtime remains *O*(*nk*), when a linear sampling of quintets is used.

## 4 Theoretical Results

In this section, we provide the main theoretical results, starting with a series of lemmas and theorems that will be used in the proof of statistical consistency of QR-STAR in Theorem 2. Throughout this paper, we assume that discordance between species trees and gene trees is solely due to ILS. In establishing statistical consistency, we assume that input gene trees are true gene trees and, thus, have no gene tree estimation error. If not otherwise specified, all trees are assumed to be fully resolved (i.e., binary). Due to space constraints, most of the proofs are provided in Appendix B. We begin with some definitions and key observations.

### Definition 1 (Path length parameter)

*Let R be an MSC model species tree. Let f*(*R*) *be the length of the shortest internal branch of R and g*(*R*) *be the length of the longest internal path (i*.*e*., *a path formed from only the internal branches) of R. We define the path length parameter of R as*

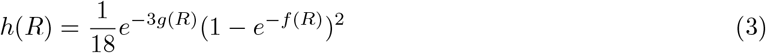

Note that 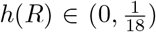 since exp(−*x*) ∈ (0, 1) for all *x >* 0 and the branch lengths have positive values. The formula for Eq. 3 is derived from the proof of Lemma 2 in Appendix B.

### Lemma 1.

*Let R be an MSC model species tree with n* ≥ 5 *leaves and q be an arbitrary set of 5 leaves from ℒL*(*R*). *Then h*(*R*|_*q*_) ≥ *h*(*R*) *where R*|_*q*_ *is the rooted tree R restricted to taxa in set q*.

### Lemma 2.

*Let R be an MSC model species tree with 5 leaves and internal branch lengths x, y, and z. Let* 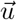 *be the probability distribution that R defines on the unrooted 5-taxon gene tree topologies. If* 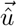 *is an estimate of* 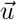 *such that given ϵ >* 0, *we have* |*û*_*i*_ − *u*_*i*_| *< ϵ for all* 1 ≤ *i* ≤ 15, *then the following inequality holds:*

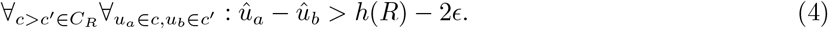

### Definition 2.

*For a 5-taxon rooted tree R, we define I*_*R*_ *as the set of ordered pairs* (*i, j*), 1 ≤ *i ≠j* ≤ 15, *corresponding to inequalities in the form u*_*i*_ *> u*_*j*_ *defined according to the partial order of R. The inequalities that are a result of transitivity (i*.*e. u*_*i*_ *> u*_*j*_ *and u*_*j*_ *> u*_*k*_ *implies u*_*i*_ *> u*_*k*_*) are not included in I*_*R*_.

### Definition 3.

*Let V* (*R, R*^*′*^) *be the set of violated inequalities of two rooted 5-taxon trees R and R*^*′*^, *i*.*e*., *all pairs* {*i, j*} *such that* (*i, j*) ∈ *I*_*R*_ *and* (*j, i*) ∈ *I*_*R*′_.

Figure 3.a shows an example of *V* (*R, R*^*′*^) computed for caterpillar trees and Figure 3.b is a heatmap showing the function |*V* (*R, R*^*′*^)| computed for the seven possible rootings of an unrooted quintet tree. The set *V* (*R, R*^*′*^) can be easily computed from *I*_*R*_ and *I*_*R*′_ for all pairs of rooted 5-taxon trees, and *I*_*R*_ is derived from the ADR theory for all 105 5-taxon rooted trees in the Supplementary Materials, Sec. S2 in [31].

**Fig. 3:**
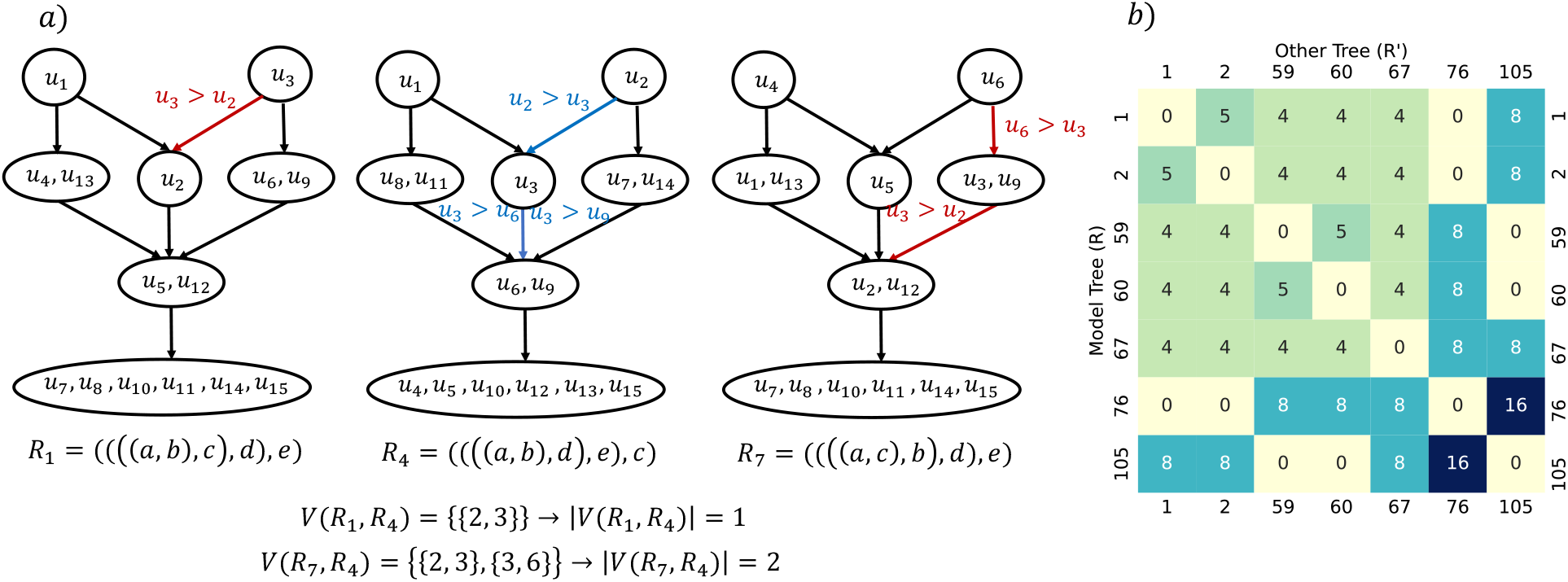
Conflicting inequality penalty terms between rooted 5-taxon species trees. a) Set of violated inequality penalty terms in the partial orders of *R*_1_ and *R*_7_ with respect to *R*_4_, which are all caterpillar trees. The red edges show violations of inequalities in tree *R*_4_, highlighted in blue. b) Heatmap showing the number of pairwise violated penalty terms (function |*V* (*R, R*^*′*^)|) of seven possible rooted trees having unrooted topology with bipartitions *ab*|*cde* and *abc*|*de*. The dark colors indicate more violations, and the lightest color corresponds to no violations (|*V* (*R, R*^*′*^)| = 0).

### Lemma 3.

*(a) For 5-taxon binary rooted trees R and R*^*′*^ *with the same rooted shape, the set V* (*R, R*^*′*^) *is always non-empty. (b) For each balanced tree B, there exist two caterpillar trees C*_1_ *and C*_2_ *such that V* (*B, C*_*i*_) = ∅.

### 4.1 Statistical Consistency

In this section, we establish statistical consistency for QR-STAR under the MSC and provide the sufficient condition for a set of sampled quintets to lead to consistency. That is, we prove that as the number of input true gene trees increases, the probability that QR-STAR and its variants correctly root the given unrooted species tree converges to 1. We first prove statistical consistency for QR-STAR when the model tree has only five taxa in Theorem 1 and then extend the proofs to trees with arbitrary numbers of taxa in Theorem 2. The main idea of the proof of consistency for 5-taxon trees is that we show as the number of input gene trees increases, the cost of the true rooted tree becomes arbitrarily close to zero, but the cost of any other rooted tree is bounded away from zero, where the bound depends on the path length parameter of the model tree, *h*(*R*). Our proofs imply sample complexity bounds. For lack of space, we leave them for the journal version.

#### Lemma 4.

*Let R be an MSC model species tree with n* ≥ 5 *leaves and Q be a set of quintets of taxa from ℒ* (*R*). *Given δ >* 0 *and k >* 0 *unrooted gene tree topologies, the following inequality holds, where* 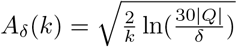

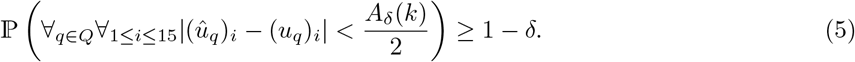

Setting 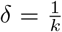 in Eq. 5, we get 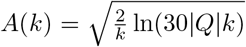, which is the bound that is used for determining the rooted shape of each quintet in the first step of QR-STAR as well as the proofs of statistical consistency. When *R* has only five taxa, *A*(*k*) becomes 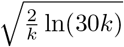, as *Q* can only contain one quintet.

#### Lemma 5 (Correct determination of rooted shape)

*Let R be a 5-taxon model species tree and* 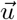 *be the probability distribution that it defines on the unrooted 5-taxon gene tree topologies. There is an integer k >* 0 *such that if we are given at least k unrooted gene trees drawn i*.*i*.*d. from the distribution* 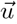, *the first step of QR-STAR will correctly determine the rooted shape of R with probability at least* 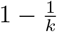.

#### Lemma 6 (Upper bound on the cost of the model tree)

*Let R be a 5-taxon model species tree and* 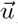 *be the probability distribution that it defines on the unrooted 5-taxon gene tree topologies. There is an integer k >* 0 *such that if we are given at least k unrooted gene trees drawn i*.*i*.*d. from distribution* 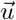, *then* 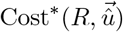 *is less than* 31*α*_max_*A*(*k*) *with probability at least* 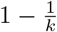.

#### Theorem 1 (Statistical Consistency of QR-STAR for 5-taxon trees)

*Let R be a 5-taxon model species tree and* 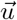 *be the distribution that it defines on the unrooted 5-taxon gene tree topologies. Given a set 𝒢 of unrooted true quintet gene trees drawn i*.*i*.*d. from* 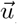, *QR-STAR is a statistically consistent estimator of R under the MSC*.

*Proof*. According to Lemma 6, when *k* is large enough so that 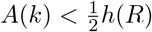, if 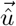 is the distribution estimated from 𝒢, then 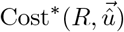 is at most 31*α*_max_*A*(*k*) with probability at least 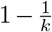. We now prove that for every other rooted 5-taxon tree *R*^*′*^, 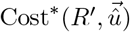 is bounded away from zero. Note that according to Lemma 3, for every rooted 5-taxon tree *R*^*′*^ ≠ *R* with the same rooted shape as *R*, we have *V* (*R, R*^*′*^)≠ ∅ and therefore, there exists 1 ≤ *x*≠ *y* ≤ 15 such that (*x, y*) ∈ *I*_*R*_, (*y, x*) ∈ *I*_*R*′_.

Let ℰ be the event that 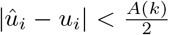 for all 1 ≤ *i* ≤ 15. According to Lemma 2, when ℰ holds, then *û*_*x*_ − *û*_*y*_ *> h*(*R*) − *A*(*k*) as (*x, y*) ∈ *I*_*R*_. However, since (*y, x*) ∈ *I*_*R*′_, an inequality penalty term in the form of max(0, *û*_*x*_ − *û*_*y*_) is added to 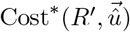 when *R*^*′*^ has the same shape as *R*. Moreover, according to Lemma 5, when ℰ holds and 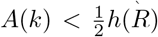, the first step of QR-STAR correctly determines the rooted shape of *R*. Therefore, if *R*^*′*^ has a different rooted shape than *R*, then the penalty 𝟙|*S*(*R*^*′*^) ≠ *Ŝ*(*û*)| becomes 1 and a positive cost *C* is added to the cost of *R*^*′*^. Therefore, both cases (i.e. whether *R*^*′*^ has a different topology from *R* or not) lead to a positive penalty in the cost function for *R*^*′*^, that is bounded away from zero. Hence,

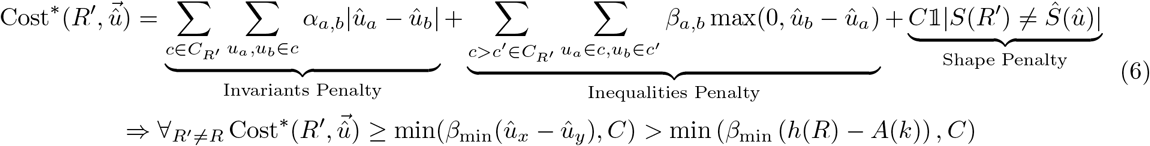

Eq. 6 defines a lower bound for the cost of any tree other than the true rooted tree and Lemma 6 gives an upper bound for the cost of the true tree, both with respect to the estimated quintet distribution 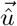. Therefore, when *k* is large enough so that

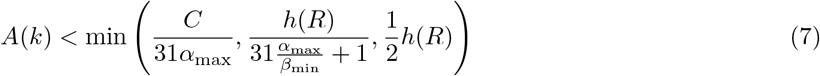

we will have

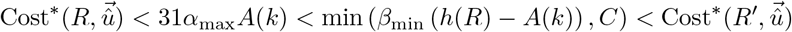

which means that the cost of the true rooted tree will be less than the cost of any other rooted tree on the same leafset with probability at least 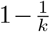, when *k* is sufficiently large. Precisely, 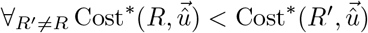 when Eq. 7 holds, where *β*_min_, *C, h*(*R*) *>* 0 and *α*_max_ ≥ 0 are constants. As a result, QR-STAR will return the true rooted species tree topology with probability converging to 1 as the number of gene trees grows large, proving the statistical consistency for 5-leaf trees. □

#### Remark 1.

Note that when *α*_max_ = 0, meaning that the invariant penalty terms are removed from the cost function, the cost of the true tree would become exactly zero according to the proof of Lemma 6, and the cost of any other tree would be positive when *k* is large enough so that the conditions of Theorem 1 hold. Hence in this case, the condition in Eq. 7 will reduce to 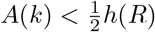.

#### Remark 2.

Note that Lemma 3(a) holds for *all* pairs of 5-taxon rooted trees with the same rooted shape and with different permutations of the leaf-labeling, regardless of whether they have the same unrooted topology or not. Due to this property, it is possible to differentiate all pairs of 5-taxon rooted trees in a statistically consistent manner with the cost function of QR-STAR without prior knowledge about the unrooted tree topology, and hence Theorem 1 does not assume that the unrooted topology is given as input.

The next lemma and theorem extend the proof of statistical consistency to trees with *n >* 5 taxa. Linear encodings of *T* are defined in Appendix A.

#### Lemma 7 (Identifiability of the root from the linear encoding)

*Let R and R*^*′*^ *be rooted trees with unrooted topology T and distinct roots. Let Q*_*LE*_(*T*) *be the set of quintets of leaves in a linear encoding of T. There is at least one quintet of taxa q* ∈ *Q*_*LE*_(*T*) *so that R*_|*q*_ *and* 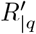 *have different rooted topologies*.

Lemma 7 states that no two distinct rooted trees with topology *T* induce the same set of rooted quintet trees on quintets of taxa in set *Q*_*LE*_(*T*). Clearly, the same is true for any superset *Q* such that *Q*_*LE*_(*T*) ⊆ *Q*, including the set *Q*_5_ of all quintets of taxa on the leafset of *T*. There might also be other quintet sets that are not a superset of *Q*_*LE*_(*T*), but have the property that no two rooted versions of *T* define the same set of rooted quintets on their elements. We generalize the proof of consistency to all set of sampled quintets with this property.

#### Definition 4.

*Let T be an unrooted tree and Q be a set of quintets of taxa from ℒ* (*T*). *We say Q is “root-identifying” if every rooted tree R with topology T is identifiable from T and the set of rooted quintet trees in* {*R*|_*q*_ : *q* ∈ *Q*}, *i*.*e*., *no two rooted trees with topology T induce the same set of rooted quintet trees on Q*.

#### Theorem 2 (Statistical Consistency of QR-STAR)

*Let R be an MSC model species tree with n* ≥5 *leaves and let T denote its unrooted topology. Given T and a set 𝒢 of unrooted true gene trees on the leafset* ℒ (*T*), *QR-STAR is a statistically consistent estimator of the rooted version of T under the MSC, if the set of sampled quintets Q is root-identifying*.

*Proof*. QR-STAR computes the score of each rooted tree *R* with topology *T* as 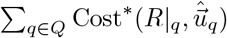. Let ℰ_*q*_ be the event that 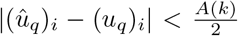 for all 1 ≤ *i* ≤ 15(for a quintet *q* ∈ *Q*. According to Theorem 1, when ℰ_*q*_ holds and *k* is large enough so that 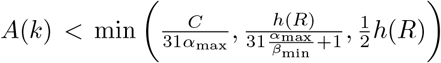 (that guarantees 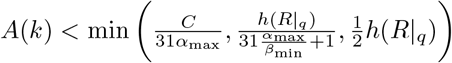 according to Lemma 1), 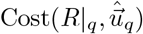 will be less than the cost of each of the six alternative rooted trees on the five taxa in *q* with high probability, as *R*|_*q*_ is the *true* rooting of the unrooted quintet tree *T* |_*q*_.

According to Lemma 4, ℰ_*q*_ simultaneously holds for all *q* ∈ *Q* with probability at least 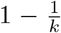. In this event, for every other rooted tree *R*^*′*^≠ *R* and each *q* ∈ *Q*, we have 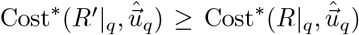, according to Theorem 1. This means that for every rooted tree *R*^*′*^ with topology *T*, 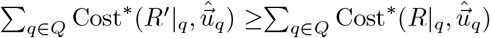. Also, according to Definition 4, *Q* has the property that no other rooted tree *R*^*′*^ induces the exact same set of rooted quintet trees as *R* on *Q*. Hence, there exists a *q*^*^ ∈ *Q* such that 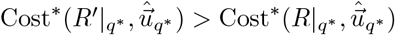, as the cost of *R*|_*q*_* is strictly less than the cost of each of the six alternative rooted trees on *q*^*^, when the conditions in Theorem 1 hold. Therefore, the function 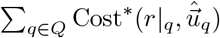 obtains its *unique* minimum for the rooted tree *R*. As a result, QR-STAR returns the true rooted topology of *T* with probability converging to 1 as the number of input gene trees increases and this proves the statistical consistency for trees with an arbitrary number of taxa.

## 5 Experimental Study

We performed an experimental study on simulated datasets to explore the parameter space of QR-STAR on a training dataset, and then compared its accuracy to QR on a test dataset. We used the 101-taxon simulated datasets from [35] as our training data, which had model conditions characterized by four levels of gene tree estimation error (GTEE) ranging from 0.23 to 0.55 (measured in terms of normalized Robinson-Foulds (RF) [27] distance between true and estimated gene trees) for 1000 genes. The normalized RF distance between the model species tree and true gene trees (denoted average distance, or AD) in this dataset was 0.46, which indicates moderate ILS. We used a set of 201-taxon simulated datasets from [20] as our test data; these are characterized by two different speciation rates and three tree heights (thus six tree shapes), and three number of genes for each tree shape. The AD levels for this dataset for 1000 genes (given in Table D1) ranged from 0.09 (for the 10M, 1e-07 condition) to 0.69 (for the 500K, 1e-06 condition). The GTEE levels on the test data varied from 0.22 (for low ILS) to 0.49 (for high ILS). The number of replicates for each model condition for both the training and test datasets was 50.

We measured the error in the rooted species tree in terms of average normalized clade distance (nCD) [31], which is an extension of RF error for rooted trees. For our training experiment, we only rooted the true species tree topology to directly observe the rooting error. In our test experiments, we rooted both the model species tree and estimated species tree, as produced by ASTRAL, using both true and estimated gene trees (which were estimated using FastTree [24]). Additional information about the simulation study, datasets, and software commands are provided in Supplementary Materials Section D.

In our training experiments, we explored the impact of the shape coefficient *C* and the ratio 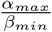 (that describes the relative impact of invariants and inequalities) on the accuracy of QR-STAR. Results for the training experiments are provided in Supplementary Materials Sec. E.1 and show that there are wide ranges of settings for the algorithmic parameters that provide the best accuracy. We used these training results and theoretical considerations related to sample complexity of QR-STAR to set the algorithmic parameters to *C* =1e-02 and 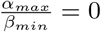.

Figure 4 shows a comparison between QR and QR-STAR in terms of rooting error for rooting the model species tree topology using estimated gene trees (GTEE values for each condition are provided in Table D1) on the test datasets. Increasing the ILS level (by reducing tree height) decreases the rooting error, and increasing the number of genes also generally reduces rooting error (although much less under the lowest ILS level where tree height is 10M). To understand the impact of ILS in Fig. 4, note that the true species tree is being rooted and so ILS will not impact the accuracy of the unrooted species tree topology. However, the level of ILS impacts information about rooting location, which comes from the distribution of gene tree topologies (see Sec. 6 for a discussion on this trend).

**Fig. 4:**
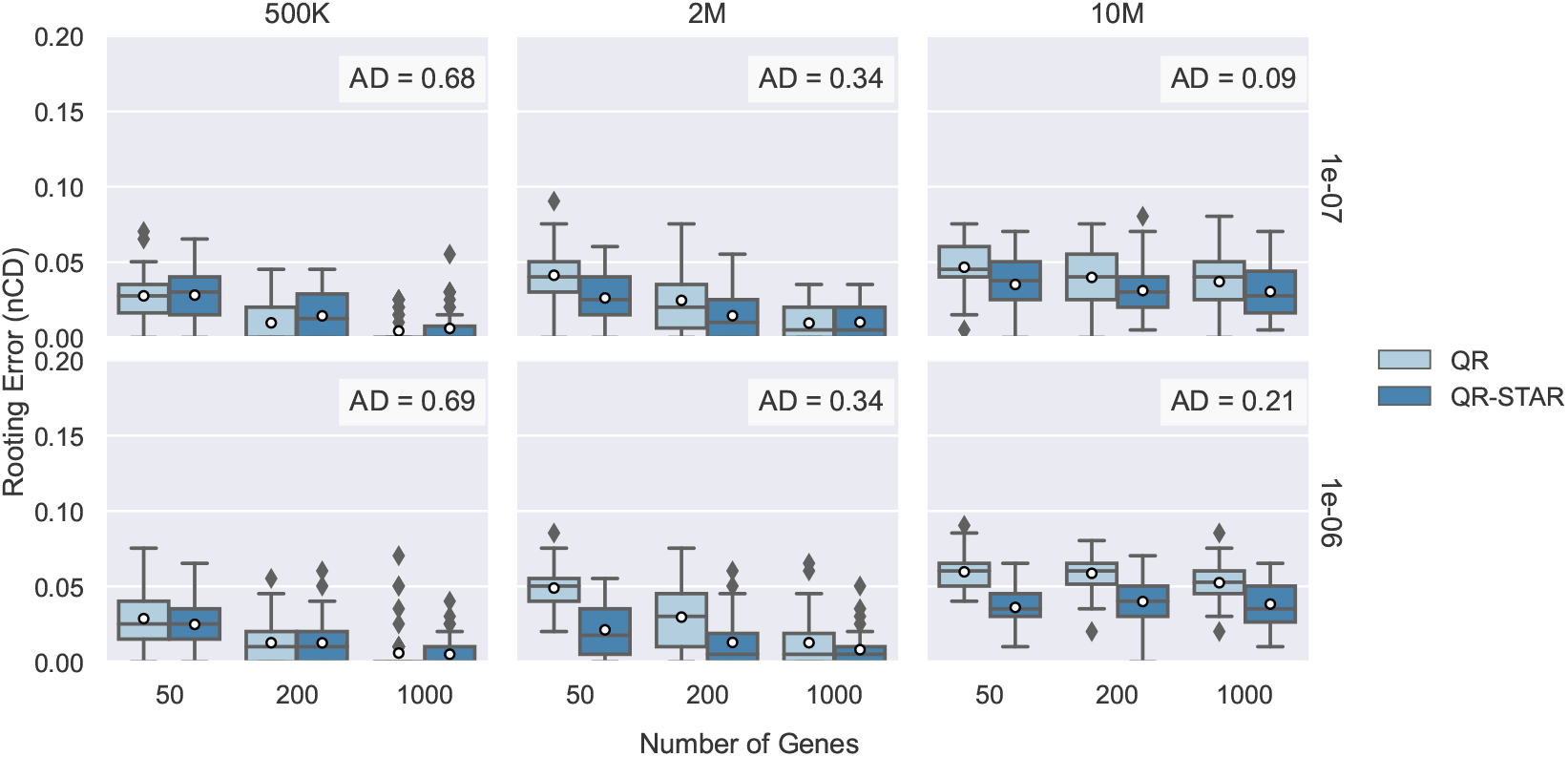
Rooting the model species tree with estimated gene trees on S200 datasets. Comparison between QR and QR-STAR in terms of rooting error (nCD) for rooting the true unrooted species tree topology using estimated gene trees (GTEE levels vary from 0.22 (for low ILS) to 0.49 (for high ILS)) on the 201-taxon datasets from [20] with 50 replicates in each model condition. The columns show tree height (500K for high ILS, 2M for moderate ILS, and 10M for low ILS), and the rows show speciation rate (1e-06 or 1e-07).

A comparison between QR and QR-STAR shows that QR-STAR generally matched or improved on QR; the only exception was for the high ILS conditions, where the two methods were very close but with perhaps a small advantage to QR. On these high ILS conditions, however, GTEE is also large, and QR-STAR is more accurate than QR when used with true gene trees, even under high ILS (see Fig. E3). Hence, the issue is likely to be high GTEE rather than high ILS, suggesting that QR-STAR is slightly more affected by GTEE compared to QR.

Figures E3, E4 and E5 show additional results on the test dataset when rooting species trees estimated using ASTRAL, with true or estimated gene trees. The relative accuracy of QR and QR-STAR in these figures are the same as seen in Figure 4, with QR-STAR performing better in almost all model conditions, but the error in the final rooted tree is impacted by both species tree estimation error and rooting error, and it is generally higher under high ILS settings.

## 6 Discussion and Concluding Remarks

In this work we presented QR-STAR, a polynomial time statistically consistent method for rooting species trees under the multispecies coalescent model. QR-STAR is an extension to QR, a method for rooting species trees introduced in [31]. QR-STAR differs from QR in that it has an additional step for determining the topological shape of each unrooted quintet selected in the QR algorithm, and incorporates the knowledge of this shape in its cost function, alongside the invariants and inequalities previously used in QR. We also showed that the statistical consistency for QR-STAR holds for a larger family of optimization problems based on cost functions and sampling methods.

To the best of our knowledge, this is the first work that established the statistical consistency of any method for rooting species trees under a model that incorporates gene tree heterogeneity. It remains to be investigated whether other rooting methods can also be proven statistically consistent under models of gene evolution inside species trees, such as the MSC or models of GDL. For example, STRIDE [4] and DISCO+QR [34] are methods that have been developed for rooting species trees from gene family trees, where genes evolve under gene duplication and loss (GDL); however, it is not known whether these methods are statistically consistent under any GDL model.

This study suggests several directions for future research. For example, we proved statistical consistency for one class of cost functions, which was a linear combination of the invariant, inequality and shape penalty terms; however, cost functions in other forms could also be explored and proven statistically consistent. Equation 7 suggests that the sample complexity of QR-STAR depends on the function *h*(*R*), which is based on both the length of the shortest branch and the longest path in the model tree. This suggests that having very short or very long branches can both confound rooting under ILS, which is also suggested in previous studies [2,1]. This is unlike what is known for species tree estimation methods such as ASTRAL, where the sample complexity is only affected by the shortest branch of the model tree [29,3], and trees with long branches are easier to estimate.

Another theoretical direction is the construction of the rooted species tree directly from the unrooted gene trees. As explained in Remark 2, the proof of consistency of QR-STAR for 5-taxon trees does not depend upon the knowledge of the unrooted tree topology; this suggests that it is possible to estimate the rooted topology of the species tree in a statistically consistency manner *directly* from unrooted gene tree topologies. Future work could focus on developing statistically consistent methods for this problem, which is significantly harder than the problem of rooting a given tree.

There are also directions for improving empirical results. An important consideration in designing a good cost function is its empirical performance, as many cost functions can lead to statistical consistency but may not provide accurate estimations of the rooted tree in practice (see Figures E1 and E2). One potential direction is to incorporate estimated branch lengths, whether of the gene trees or the unrooted species tree, into the rooting procedure.

Finally, one of the perhaps interesting results of the experimental study in this paper shows that rooting with QR and QR-STAR is easier under higher levels of discordance due to ILS, and becomes more difficult as the ILS level decreases. An explanation for this could be that with less discordance, it is likely that many gene trees that have low probability of appearing will not appear in the input. In this case, some estimates of quintet probabilities would become zero, and it may not be possible to differentiate some of the rooted quintets using the inequality and invariants derived from the ADR theory. In the extreme case, when there is no discordance, there will be only one quintet gene tree with non-zero probability, and the identifiability theorem in [2] would not hold and it becomes impossible to find the root. This as well is unlike what is known for species tree estimation methods, as they generally show the reverse trend: the accuracy of the species tree methods decrease under higher levels of ILS [19,20,22]. This suggests that an interesting trade-off could exist between the level of discordance and its impact on species tree estimation and rooting. However, the results in Figures E4 and E5 suggest that the error in the final rooted species tree might be dominated by the species tree estimation error, especially when the number of genes is small or the ILS level is high, and so the overall error in the rooted species tree would increase as the ILS levels increase.

## Acknowledgments

SR was supported by NSF grants DMS-1902892, DMS1916378 and DMS-2023239 (TRIPODS Phase II), as well as a Vilas Associates Award. TW was supported by the Grainger Foundation. SR thanks Cécile Ané and her group for helpful discussions. YT thanks Mohammed El-Kebir for helpful suggestions on an earlier version of this work. The authors thank the anonymous reviewers for their feedback.

## Code and Data Availability

QR-STAR is available at https://github.com/ytabatabaee/Quintet-Rooting. The scripts and data used in this study are available at https://github.com/ytabatabaee/QR-STAR-paper.

## Supplementary Materials

### A Quintet Sampling in QR

The set *Q* of quintets in the QR algorithm can be selected in different ways, and here we consider different sampling strategies that lead to statistical consistency. These sampling strategies differ significantly in the number of quintets they sample and therefore their runtime. A straightforward sampling strategy is to use all *Θ*(*n*^5^) quintets, which was the main approach used in [31]. Additionally, an *O*(*n*) sampling method called “Linear Encoding” was proposed, where a set *Q*_*LE*_ was selected in the following way: each edge *e* of *T* that is adjacent to a leaf *x* partitions the set of taxa into three sets {*x*}, *A, B*, and they formed a quintet *q*_*e*_ by selecting {*x*} and at least one leaf from each of the sets *A* and *B*, with the other two leaves of *q*_*e*_ being randomly selected. Similarly, each internal edge *e* partitions the set of taxa into four subsets *A*_1_, *A*_2_, *B*_1_ and *B*_2_, and a quintet *q*_*e*_ could be formed by picking at least one taxon from each of these four sets, with the final taxon being arbitrarily selected. Since a tree with *n* leaves has 2*n*− 3 edges, |*Q*_*LE*_| = 2*n*− 3 and therefore the preprocessing step of QR has a runtime of *O*(*nk*) when the sampling method used is linear encoding. Note also that *Q*_*LE*_(*T*) may not be unique for trees with *n >* 5 taxa.

### B Proofs

#### B.1 Proof of Lemma 1

*Proof*. Every internal path of *R*|_*q*_ is also an internal path in *R*, and therefore *g*(*R*|_*q*_) ≤ *g*(*R*). Also, every branch in *R*|_*q*_ is formed from one or more branches in *R*, so the shortest branch in *R*|_*q*_ should be at least as long as the shortest branch in *R*, and therefore *f*(*R*|_*q*_) ≥ *f*(*R*). Hence,

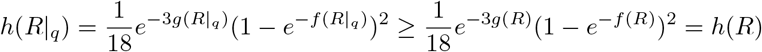

□

#### B.2 Proof of Lemma 2

*Proof*. Let *X* = *e*^−*x*^, *Y* = *e*^−*y*^ and *Z* = *e*^−*z*^. According to the explicit formulas for the probability distribution of unrooted gene trees 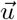 under a 5-taxon model species tree provided in Appendix B of [2], the exact value of each *u*_*i*_ can be expressed as a polynomial with variables *X, Y* and *Z*. We show that the statement of the lemma holds for all pairs *u*_*a*_, *u*_*b*_ from different equivalence classes, for each tree category. For each category, we only show that the lemma holds for one example tree with that rooted shape (trees in Table 1 in [31]), as the rest of the trees have 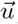 distributions that are only permutations of the distributions of these three example trees (see Supplementary Materials Sec. S2 in [31]) and the explicit formulas remain the same. The following equations can be derived using elementary algebraic arguments, and the fact that *X, Y, Z*, ∈ (0, 1).

##### Caterpillar Trees

For a caterpillar tree *R* = ((((*a, b*) : *x, c*) : *y, d*) : *z, e*) with *C*_*R*_ = {*c*_1_ : {*u*_1_}, *c*_2_ : {*u*_3_}, *c*_3_ : {*u*_2_}, *c*_4_ : {*u*_4_, *u*_13_}, *c*_5_ : {*u*_6_, *u*_9_}, *c*_6_ : {*u*_5_, *u*_12_}, *c*_7_ : {*u*_7_, *u*_8_, *u*_10_, *u*_11_, *u*_14_, *u*_15_}} we have *c*_1_ *> c*_3_, *c*_4_ *> c*_6_ *> c*_7_ and *c*_2_ *> c*_3_, *c*_5_ *> c*_6_ *> c*_7_. Therefore,

– *u*_*a*_ ∈ *c*_1_, *u*_*b*_ ∈ *c*_3_

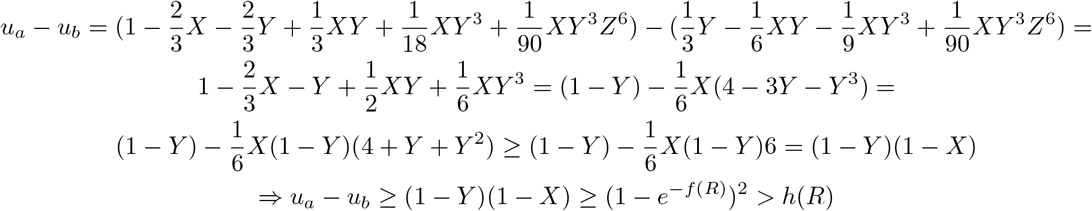

where (1 − *Y*)(1 − *X*) ≥ (1 − *e*^−*f*(*R*)^)^2^ follows from the fact that (1 − *X*) = 1 − *e*^−*x*^ ≥ 1 − *e*^−*f*(*R*)^ as *x* ≥ *f*(*R*), and same is true for *y*.
– *u*_*a*_ ∈ *c*_1_, *u*_*b*_ ∈ *c*_4_

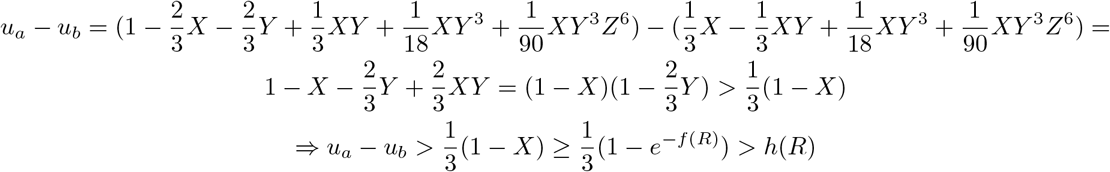
– *u*_*a*_ ∈ *c*_2_, *u*_*b*_ ∈ *c*_3_

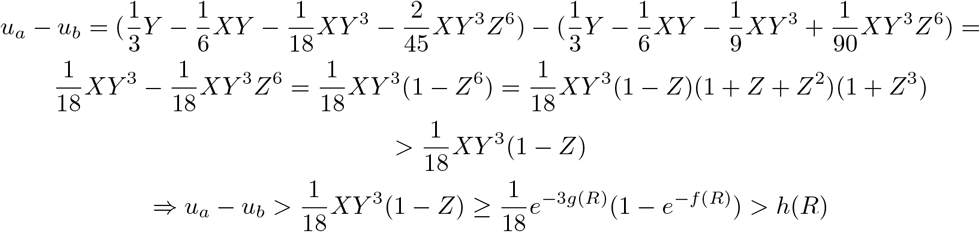

where *XY* ^3^ ≥ *e*^−3*g*(*R*)^ follows from the fact that *XY* = *e*^−*x*^*e*^−*y*^ = *e*^−(*x*+*y*)^ ≥ *e*^−*g*(*R*)^ as *x* + *y* correspond to a (sub)-length of an internal path in *R* and hence *x* + *y* ≤ *g*(*R*) and *Y* ^2^ ≥ *e*^−2*g*(*R*)^ as *y* ≤ *g*(*R*).
– *u*_*a*_ ∈ *c*_2_, *u*_*b*_ ∈ *c*_5_

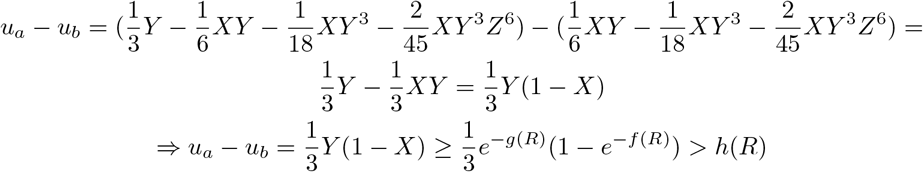
– *u*_*a*_ ∈ *c*_3_, *u*_*b*_ ∈ *c*_6_

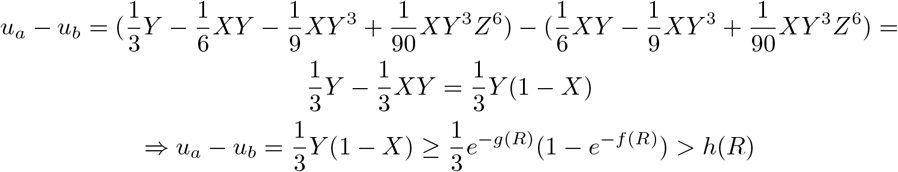
– *u*_*a*_ ∈ *c*_4_, *u*_*b*_ ∈ *c*_6_

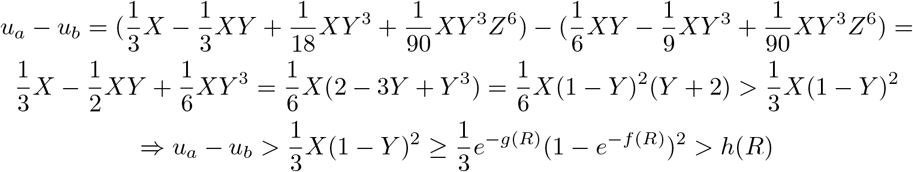
– *u*_*a*_ ∈ *c*_5_, *u*_*b*_ ∈ *c*_6_

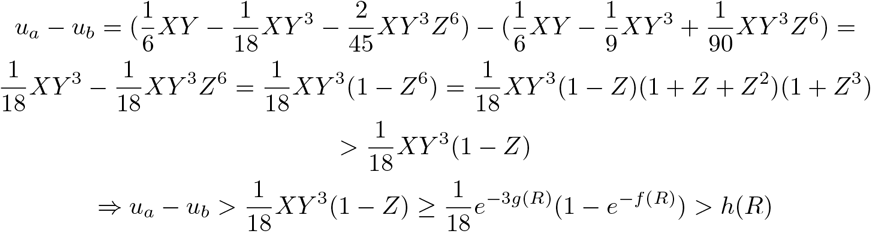
– *u*_*a*_ ∈ *c*_6_, *u*_*b*_ ∈ *c*_7_

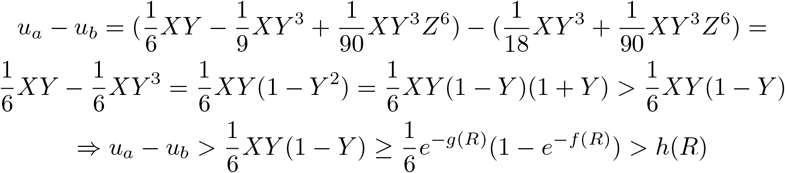

##### Balanced Trees

For a balanced model species tree *R* = (((*a, b*) : *x, c*) : *y*, (*d, e*) : *z*), with *C*_*R*_ = {*c*_1_ : {*u*_1_}, *c*_2_ : {*u*_4_, *u*_13_}, *c*_3_ : {*u*_2_, *u*_3_}, *c*_4_ : {*u*_5_, *u*_6_, *u*_9_, *u*_12_}, *c*_5_ : {*u*_7_, *u*_8_, *u*_10_, *u*_11_, *u*_14_, *u*_15_}}, we have *c*_1_ *> c*_2_, *c*_3_ *> c*_4_ *> c*_5_. Therefore,

– *u*_*a*_ ∈ *c*_1_, *u*_*b*_ ∈ *c*_3_

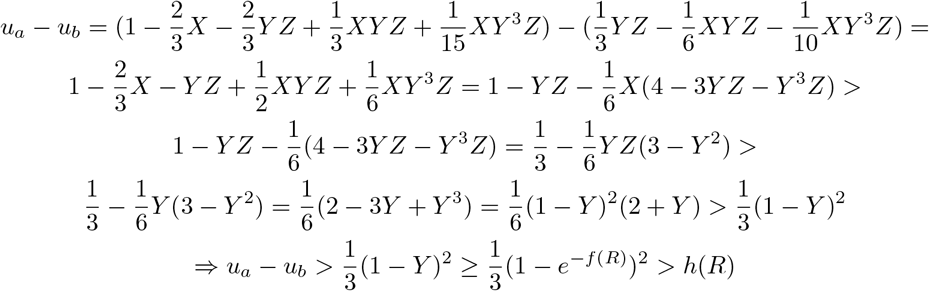
– *u*_*a*_ ∈ *c*_1_, *u*_*b*_ ∈ *c*_2_

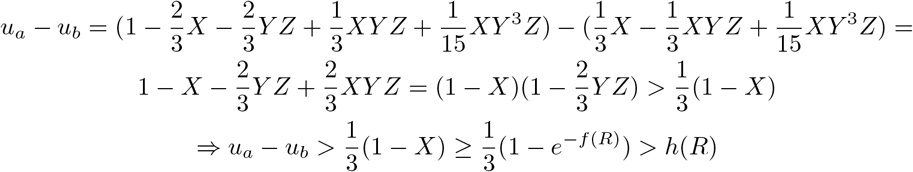
– *u*_*a*_ ∈ *c*_3_, *u*_*b*_ ∈ *c*_4_

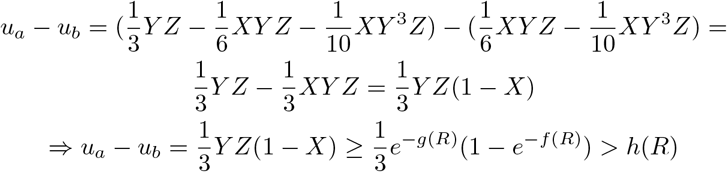
– *u*_*a*_ ∈ *c*_2_, *u*_*b*_ ∈ *c*_4_

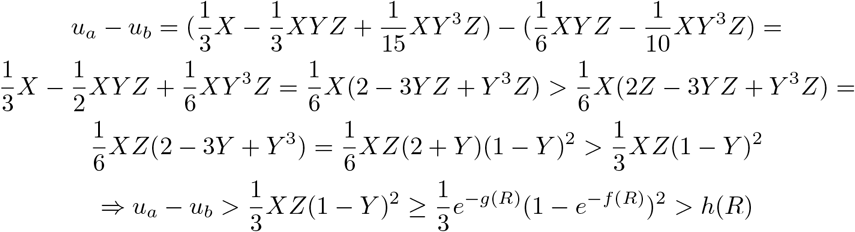
– *u*_*a*_ ∈ *c*_4_, *u*_*b*_ ∈ *c*_5_

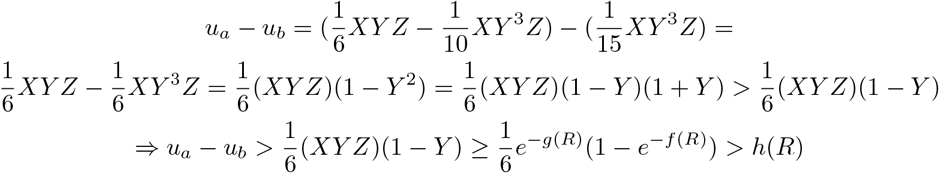

where *XY Z* ≥ *e*^−*g*(*R*)^ follows from *x* + *y* + *z* ≤ *g*(*R*).

##### Pseudo-caterpillar Trees

For a pseudo-caterpillar model species tree *R* = (((*a, b*) : *x*, (*d, e*) : *y*) : *z, c*) with *C*_*R*_ = {*c*_1_ : {*u*_1_}, *c*_2_ : {*u*_4_, *u*_13_}, *c*_3_ : {*u*_2_, *u*_3_}, *c*_4_ : {*u*_8_, *u*_11_}, *c*_5_ : {*u*_5_, *u*_6_, *u*_7_, *u*_9_, *u*_10_, *u*_12_, *u*_14_}} we have *c*_1_ *> c*_2_, *c*_3_, *c*_4_ *> c*_5_. Therefore,

– *u*_*a*_ ∈ *c*_1_, *u*_*b*_ ∈ *c*_2_

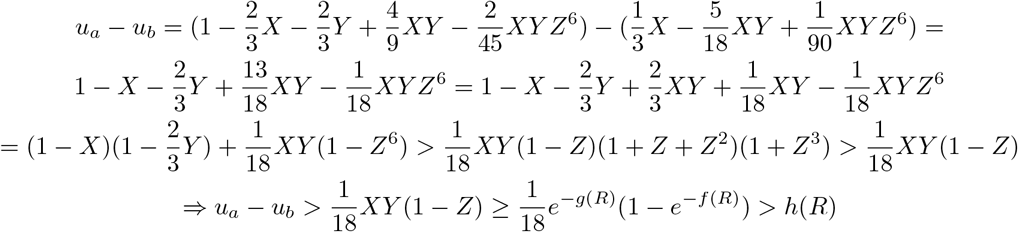
– *u*_*a*_ ∈ *c*_1_, *u*_*b*_ ∈ *c*_3_

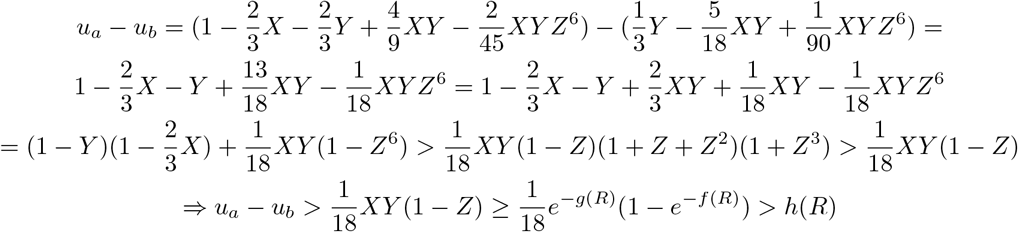
– *u*_*a*_ ∈ *c*_1_, *u*_*b*_ ∈ *c*_4_

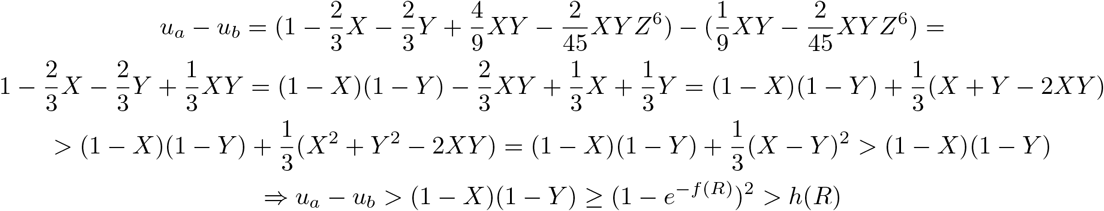
– *u*_*a*_ ∈ *c*_2_, *u*_*b*_ ∈ *c*_5_

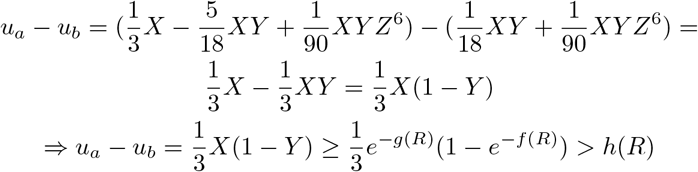
– *u*_*a*_ ∈ *c*_3_, *u*_*b*_ ∈ *c*_5_

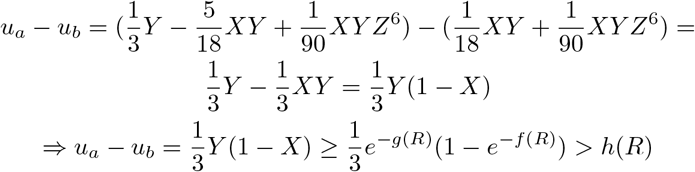
– *u*_*a*_ ∈ *c*_4_, *u*_*b*_ ∈ *c*_5_

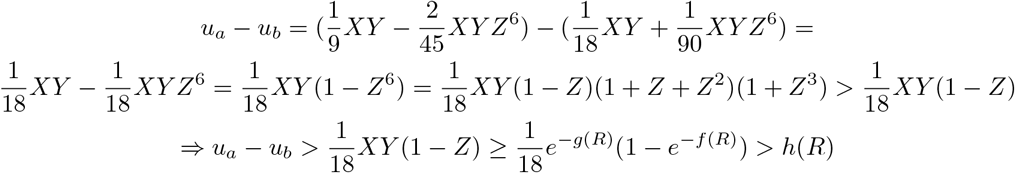

Therefore we have,

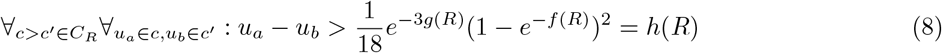

Since |*u*_*i*_ − *û*_*i*_| *< ϵ*, we have −*ϵ < û*_*i*_ − *u*_*i*_ *< ϵ* and −*ϵ < u*_*i*_ − *û*_*i*_ *< ϵ*. According to Eq. 8,

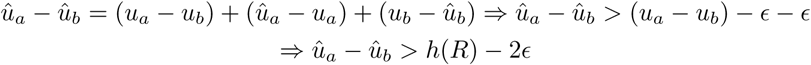

□

#### B.3 Proof of Lemma 3

*Proof*. (a) Figures F6, F8 and F7 show the function |*V* (*R, R*^*′*^)| (number of violated inequalities) for all rooted quintet tree pairs *R* and *R*^*′*^ with the same unlabeled topological shape (i.e., caterpillar, balanced and pseudo-caterpillar), computed using the invariants and inequalities derived from the ADR theory (for details on how these are computed, refer to [31], Supplementary Material Sec. 2). It is clear that, except for the numbers on the main diagonal, all other values are non-zero. Therefore, *V* (*R, R*^*′*^) is always non-empty when *R* and *R*^*′*^ have the same rooted topological shape.

(b) W.L.O.G. assume we have a particular unrooted quintet tree *T*_1_ (see Table 5 in [2]) so that its seven possible rootings are caterpillar trees *R*_1_, *R*_2_, *R*_59_, *R*_60_, pseudo-caterpillar tree *R*_67_ and balanced trees *R*_76_ and *R*_105_. Figure 3.b shows the function |*V* (*R, R*^*′*^)| for all these trees, and it is evident that for the balanced trees *R*_76_ and *R*_105_, there are two caterpillar trees (*R*_1_ and *R*_2_ for *R*_76_, and *R*_59_ and *R*_60_ for *R*_105_) for which |*V* (*R, R*^*′*^)| becomes zero. The same can be observed for trees with other unrooted topologies in Figure F9. □

#### B.4 Proof of Lemma 4

*Proof*. For an arbitrary *ϵ >* 0, we have

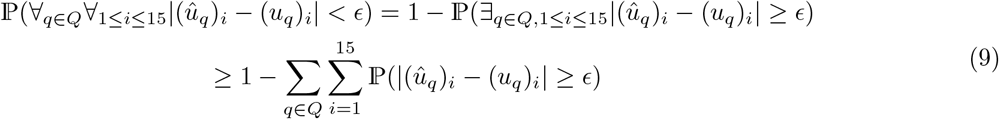

according to the union bound. Using the Hoeffding inequality [21] for each of the 15 unrooted 5-taxon tree topologies, we get

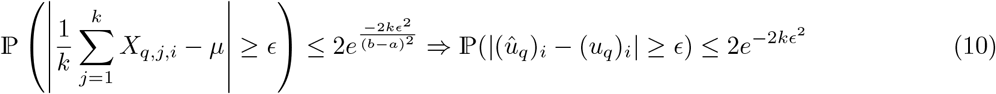

where *X*_*q,j,i*_ is a binary random variable that is 1 when the quintet gene tree *g*|_*qj*_ has the unrooted topology *T*_*i*_ and is zero otherwise, and so 0 ≤ *X*_*q,j,i*_ ≤ 1 almost surely. Substituting Eq. 10 in Eq. 9, we get

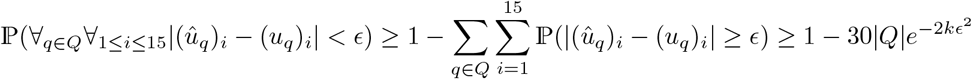

Setting 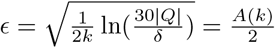 in the equation above proves the lemma

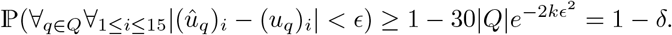

□

#### B.5 Proof of Lemma 5

*Proof*. Let ℰ be the event that 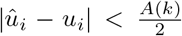 for all 1 ≤ *i* ≤ 15. Assume *k* is large enough so that 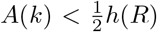. According to Lemma 4, the probability that ℰ occurs is at least 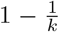. We assume that ℰ holds in the rest of this proof. In this case, according to Lemma 2, we have

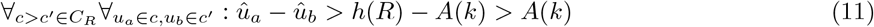

where the last inequality is a result of the assumption 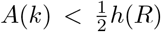. Therefore, the minimum distance between elements of any two equivalence classes that are related by an inequality in the partial order, i.e. *c > c*^*′*^ ∈ *C*_*R*_, is greater than *A*(*k*). Moreover, the maximum distance between elements inside an equivalence class is less than *A*(*k*), as according to the triangle inequality and since *u*_*a*_ = *u*_*b*_,

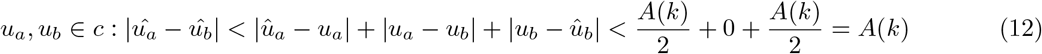

The partial order of each of the three topological shapes has a unique equivalence class with minimum probability, and for caterpillar and balanced shapes, it is followed by another unique class with second minimum probability. Since the distance between elements in different equivalence classes related by an inequality is greater than *A*(*k*), and the distance between elements inside an equivalence class is less than *A*(*k*), after sorting 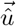 in ascending order, the elements of the equivalence class with the smallest probability appear at the beginning, followed by the elements of the second smallest class (for caterpillar and balanced shapes). Let 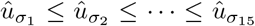 be the result of sorting 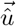 in ascending order. For a pseudo-caterpillar tree, the class with the minimum probability has 8 elements, and for caterpillar or balanced trees it has 6 elements.

– The first step of QR-STAR determines the tree shape as pseudo-caterpillar if 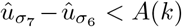 and else it will determine the shape as either caterpillar or balanced. When 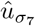 and 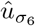 belong to different classes, their distance must be greater than *A*(*k*). Therefore, when 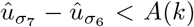 holds, 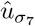 and 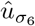 must belong to the same equivalence class, and this only happens when the model tree is a pseudo-caterpillar tree. On the other hand, if *R* is a pseudo-caterpillar tree, then 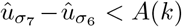 as the first eight elements in the result of the sorting belong to the same class. Therefore condition 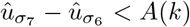 holds if and only if *R* has a pseudo-caterpillar shape and therefore QR-STAR determines the correct unlabeled shape in this case.
– If 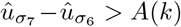, then *R* is either a balanced or a caterpillar tree. The second smallest equivalence class for both caterpillar and balanced trees is unique and has size 2 for caterpillar trees and size 4 for balanced trees. The first step of QR-STAR determines the tree shape as balanced if condition 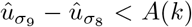 holds and else it would determine the tree shape as caterpillar. Similar to the explanation above for the case of pseudo-caterpillar trees, when 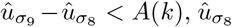 and 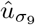 must belong to the same equivalence class and this only happens for balanced trees. Moreover, when *R* is balanced, 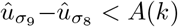. Therefore, conditions 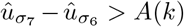 and 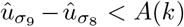 hold if and only if *R* is a balanced tree and QR-STAR correctly determines the tree shape in this case as well.
– Finally, when *R* is a caterpillar tree, 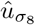 and 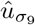 belong to different equivalence classes and therefore, 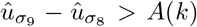and the other side can be shown similarly. Therefore, by comparing 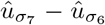 and 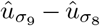 against *A*(*k*) when *k* is large enough so that 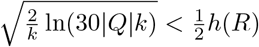, QR-STAR will correctly determine the rooted shape of the model tree with probability at least 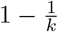.

The argument is summarized below:

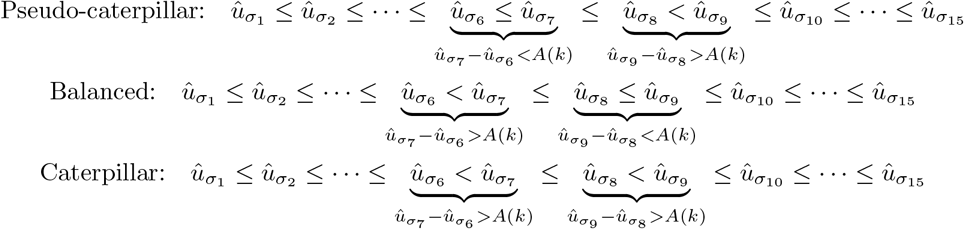

□

#### B.6 Proof of Lemma 6

*Proof*. Let ℰ be the event that 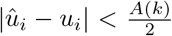 for all 1 ≤ *i* ≤ 15. According to Lemma 4, the probability that ℰ holds is at least 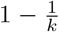. When ℰ holds, according to Lemma 2, the following inequality is true for 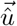 and the path length parameter *h*(*R*) of the model tree *R*

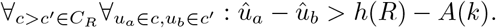

When *k* is sufficiently large so that 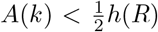, then *û*_*a*_ − *û*_*b*_ will be positive. Therefore, all inequality penalty terms which are defined as max(0, *û*_*b*_ *û*_*a*_) in 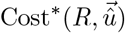 become zero, since *û*_*b*_ *û*_*a*_ is a negative term. Therefore, the total sum of the inequality penalty terms in 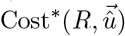 will be zero for large enough *k*. Moreover, Lemma 5 states that the topological shape of *R* can be correctly determined when 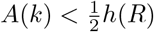 and ℰ holds, and hence the shape penalty term 𝟙|*S*(*R*)≠ *Ŝ*(*û*)| also becomes zero. Therefore, all elements of 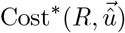 except the invariants penalty terms become zero.

According to Eq. 12, for each invariant penalty term, we have |*û*_*a*_ − *û*_*b*_| *< A*(*k*). Hence,

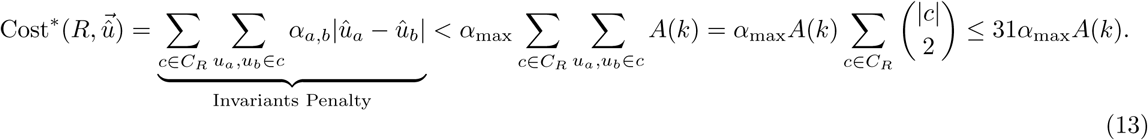

The last inequality holds since caterpillar trees have 7 equivalence classes with class sizes 1,1,1,2,2,2,6 and therefore 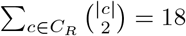, balanced trees have 5 equivalence classes with sizes 1,2,2,4,6 and 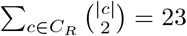 and pseudo-caterpillar trees have 5 equivalence classes with sizes 1,2,2,2,8 and 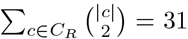. Hence, in all cases we have 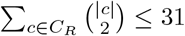 and the inequality follows. □

#### B.7 Proof of Lemma 7

*Proof*. Let *e* be the edge in *T* corresponding to the root of *R*. Let *q*(*e*) be the quintet of leaves corresponding to edge *e* in *Q*_*LE*_(*T*). It is clear that *R*_|*q*(*e*)_ is also rooted at edge *e*. The following cases can happen:

– When edge *e* is not incident to a leaf, it partitions the set of leaves of *T* into four subsets. Let *A*_1_, *A*_2_, *B*_1_ and *B*_2_ be the subsets resulting from deleting edge *e* from *T*, where *A*_1_ and *A*_2_ are adjacent to one endpoint of *e* and *B*_1_ and *B*_2_ are adjacent to the other. According to the definition of quintets in linear encoding, *q*(*e*) must have at least one leaf in each subset, which we call *a*_1_, *a*_2_, *b*_1_ and *b*_2_ respectively. For every other rooted tree *R*^*′*^ with topology *T*, the root edge *e*^*′*^ of *R*^*′*^ will fall into one of the four subsets *A*_1_, *A*_2_, *B*_1_, *B*_2_, including the edges directly adjacent to *e*. W.L.O.G. assume that *e*^*′*^ falls into *A*_1_. Then when *T* is rooted at edge *e*^*′*^ (resulting in *R*^*′*^), the leaves *a*_2_, *b*_1_ and *b*_2_ in *q*(*e*) fall into one side of the root and *a*_1_ falls into another. Therefore, the leaves *a*_1_, *a*_2_ and *b*_1_ form the rooted triplet ((*a*_2_, *b*_1_), *a*_1_). However, when *T* is rooted at *e* (producing *R*), these leaves form the rooted triplet ((*a*_1_, *a*_2_), *b*_1_), as edge *e* separates *a*_1_ and *a*_2_ from *b*_1_. Hence, in this case *R*_|*q*(*e*)_ and 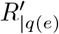 are topologically different because they induce different rooted triplets.
– When edge *e* is adjacent to a leaf *x*, it partitions the set of taxa into three subsets, where one of them contains the single node {*x*}. Let *A, B* be the two other subsets resulting from deleting the edge *e*, where *e* separates *x* from the sets *A* and *B*. According to the definition of linear encoding, *q*(*e*) must contain *x* and at least one leaf in *A* and *B* which we call *a* and *b* respectively. For every other rooted tree *R*^*′*^ with topology *T*, the root edge *e*^*′*^ of *R*^*′*^ will fall into *A* or *B* (including the edges directly adjacent to *e*). W.L.O.G. assume that it falls in *A*. Then when *T* is rooted at *e* (producing *R*), the nodes *a, b, x* form the rooted triplet ((*a, b*), *x*), but when *T* is rooted at *e*^*′*^ (producing *R*^*′*^), they form the rooted triplet ((*b, x*), *a*). Therefore, this case also leads to different rooted topologies for *R*_|*q*(*e*)_ and 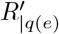

Therefore, in both cases, *R* and *R*^*′*^ restricted to quintet *q*(*e*) produce topologically different rooted quintet trees and this proves the lemma. □

### C Lack of consistency of QR

The original QR algorithm uses the following cost function to select between different rootings of a 5-taxon unrooted species tree, given the estimated quintet distribution 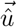:

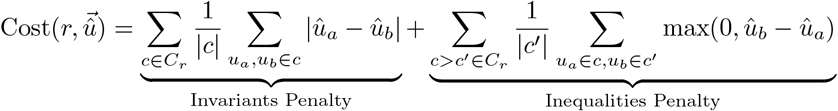

Lemma 3 shows that, for each balanced tree, there are two caterpillar trees for which the set of violated inequalities becomes empty. Consider the caterpillar tree *R*_1_ and balanced tree *R*_76_ shown in Figure 1. *Note that class c*_3_ *in R*_76_ *is the result of merging the classes c*_2_ *and c*_3_ *in R*_1_. *Moreover, class c*_4_ *in R*_76_ *is the result of merging classes c*_5_ *and c*_6_ *in R*_1_. Assume that the model tree is *R*_76_, and we have estimated 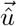 given a set of *k* unrooted quintet gene trees. We now argue informally that even as *k* increases, there is no guarantee that eventually 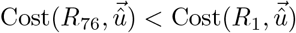 for all large enough *k*.

According to the proof of Lemma 6, as *k* increases, for the model tree *R*_76_ all inequality penalty terms in the form of max(0, *û*_*b*_ − *û*_*a*_) will converge to zero in probability. Therefore, roughly speaking, the cost of *R*_76_ eventually consists primarily of the invariant penalty terms. For the caterpillar tree *R*_1_, most of its inequality penalty terms are also a penalty term in the cost of *R*_76_, but it also has additional penalty terms between classes *c*_2_ and *c*_3_ as well as classes *c*_5_ and *c*_6_ that are merged in *R*_76_. By simplifying the penalty terms that are eventually zero with high probability or are included in the cost of both trees, we get

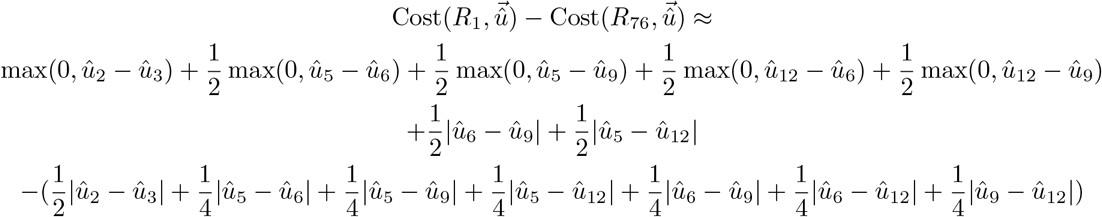

In the limit *k* → +∞, all remaining terms in that difference also go to 0 under the model tree. Hence, intuitively, there is no guarantee that *R*_76_ will be selected by QR. Based on this informal argument, we conjecture that QR is not statistically consistent.

### D Details of the Experimental Study

#### D.1 Details of the Datasets

Table D1 provides the average AD and GTEE values for the six model conditions of the test dataset (S200 dataset from [20]). The two speciation rates used in this dataset indicate whether speciation happened close to the leaves (i.e. recent speciation for 1e-06) or close to the root (i.e. deep speciation for 1e-07). The shorter tree height (500K), indicates shorter branches and therefore higher levels of ILS. Table D1 shows that the two high-ILS conditions have relatively higher levels of gene tree estimation error as well. For the training dataset (S100 dataset from [35]), the AD level for most replicates range from 0.3 to 0.6 with an average of 0.46. The mean GTEE values for the four sequence lengths is 0.23, 0.31, 0.42 and 0.55 for the 1600bp, 800bp, 400bp and 200bp sequences respectively. The speciation rate for this dataset was 1e-07.

#### D.2 Software Commands and Version Numbers

**– ASTRAL:** We used ASTRAL (v5.7.8) to estimate unrooted species trees. ASTRAL is available at https://github.com/smirarab/ASTRAL. We used the following command:

~~~
java - jar astral .5 .7 .8. jar -i <input - genes. tre > -o < output. tre >
~~~

**Table D1:** Statistics for the S200 datasets. True average discordance between the true gene trees and the model species tree (AD level) and average gene tree estimation error (GTEE) of the estimated gene trees for the six model conditions of the S200 datasets with 1000 gene trees.

| speciation rate | tree height | true discordance (AD) | GTEE |
| --- | --- | --- | --- |
| 1E-06 | 500K | 0.69 | 0.49 |
| 1E-06 | 2M | 0.34 | 0.28 |
| 1E-06 | 10M | 0.21 | 0.22 |
| 1E-07 | 500K | 0.68 | 0.46 |
| 1E-07 | 2M | 0.34 | 0.34 |
| 1E-07 | 10M | 0.09 | 0.29 |

– **QR:** We used QR (v1.2.4) to root unrooted species trees. QR is available at https://github.com/ytabatabaee/Quintet-Rooting. We used the following command:

~~~
python 3 quintet_rooting - py -t <input - tree. tre > -g <input - genes. tre > -o < output. tre >
- sm le
~~~ The `-LE` option specifies the quintet sampling method as “linear encoding”.
– **QR-STAR:** QR-STAR is available as part of the QR software package, and we ran it using the following command in the comparisons to QR, that sets *C* = 1*E* − 02 and 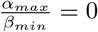:

~~~
python 3 quintet_rooting. py -t <input - tree. tre > -g <input - genes. tre > -o < output. tre >
- sm le -c STAR - abratio 0 - coef 0.01
~~~
– **GTEE, AD and Rooting (nCD) Error:** Gene tree estimation error and average discordance between model species trees and true gene was computed using a script for computing normalized RF distance written by Erin. K. Molloy available at https://github.com/ekmolloy/njmerge/blob/master/python/compare_trees.py..

Rooting error was measured in terms of average normalized clade distance (nCD) using the script available at https://github.com/ytabatabaee/Quintet-Rooting/blob/main/scripts/clade_distance.py

### E Additional Results

#### E.1 Exploring the Parameter Space in QR-STAR

We explored a range of values for the shape coefficient (parameter *C*) and the relative weight of inequalities and invariants (the ratio 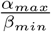) in the cost function of QR-STAR on the training dataset. Eq. 7 suggests that these two values can impact the sample complexity of QR-STAR as well. We report the proportion of the trees (from the 50 replicates in each condition) that are correctly rooted, as well as the rooting error (nCD values) for rooting the true species tree topology. Figure E1 shows the impact of shape coefficient on the accuracy of QR-STAR, where the weights of invariant and inequality penalty terms are fixed to the weights in the original cost of QR. The value of *C* varies between 0 to to 10^3^, with 0 corresponding to the cost of QR that does not guarantee consistency. For small *C* values (i.e. less than 1E-02), the accuracy of QR-STAR does not seem to be affected by the shape coefficient, but as *C* gets larger, the accuracy degrades until it reaches a stationary point again. This suggest that the shape coefficient should be kept relatively small compared to the invariant and inequality penalty weights, as they may better capture the difference between two rooted trees. Since Eq. 7 suggests that larger *C* values are theoretically preferred, on the experiments on the test dataset, we set the value of *C* as 1E-02 (the largest value before accuracy degrades).

Figure E2 shows the impact of the ratio 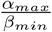 on QR-STAR. Here all *α* and *β* values are set as equal. The results suggest that when the inequalities are weighed more than the invariants (and so 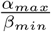 is less than 1), QR-STAR has its optimal accuracy, and the accuracy degrades when the invariants are weighed more. For both figures, the trends for different sequence lengths are similar, and the degradation in accuracy starts almost at the same point, but the accuracy is higher for longer sequence lengths, which is expected as shorter sequence lengths correspond to higher levels of GTEE. In general, these experiments show that optimal accuracy could be achieved for a wide range of parameters in QR-STAR. For experiments on the test dataset, we set *C* as 1E-02 and 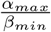 as 0 (essentially removing invariants from the cost function), although we note that the optimal values could be dataset-dependant.

**Fig. E1:**
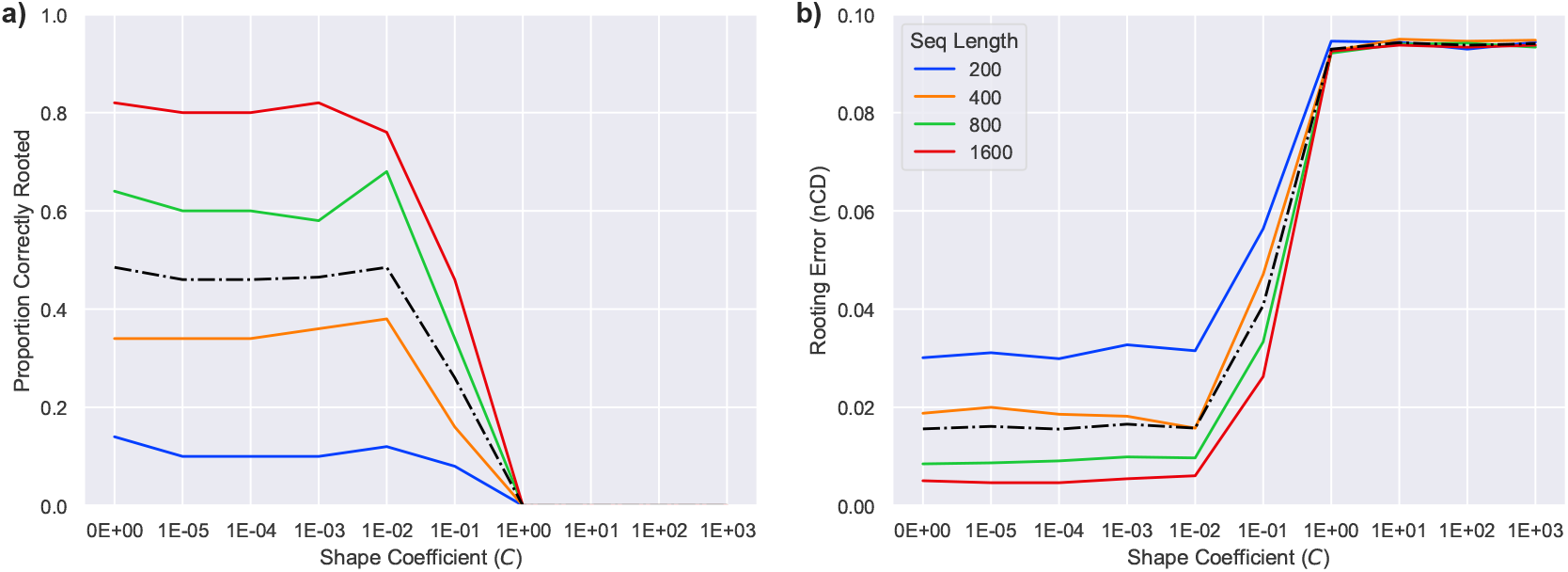
Impact of shape coefficient (*C*) on QR-STAR. a) Proportion of the trees correctly rooted and b) rooting error (nCD) are shown for the S100 dataset from [35] with 50 replicates. The number of taxa is 101, the number of genes is 1000 and the average AD level is 0.46. The sequence length used to produce estimated gene trees varies between 200bp to 1600bp. The black dashed line corresponds to the average among sequence lengths. The weights of invariant and inequality penalty terms are set as in the cost function of QR. The value of *C* varies between 10^−5^ to 10^3^ in addition to 0, that corresponds to the cost function of QR that does not guarantee consistency.

**Fig. E2:**
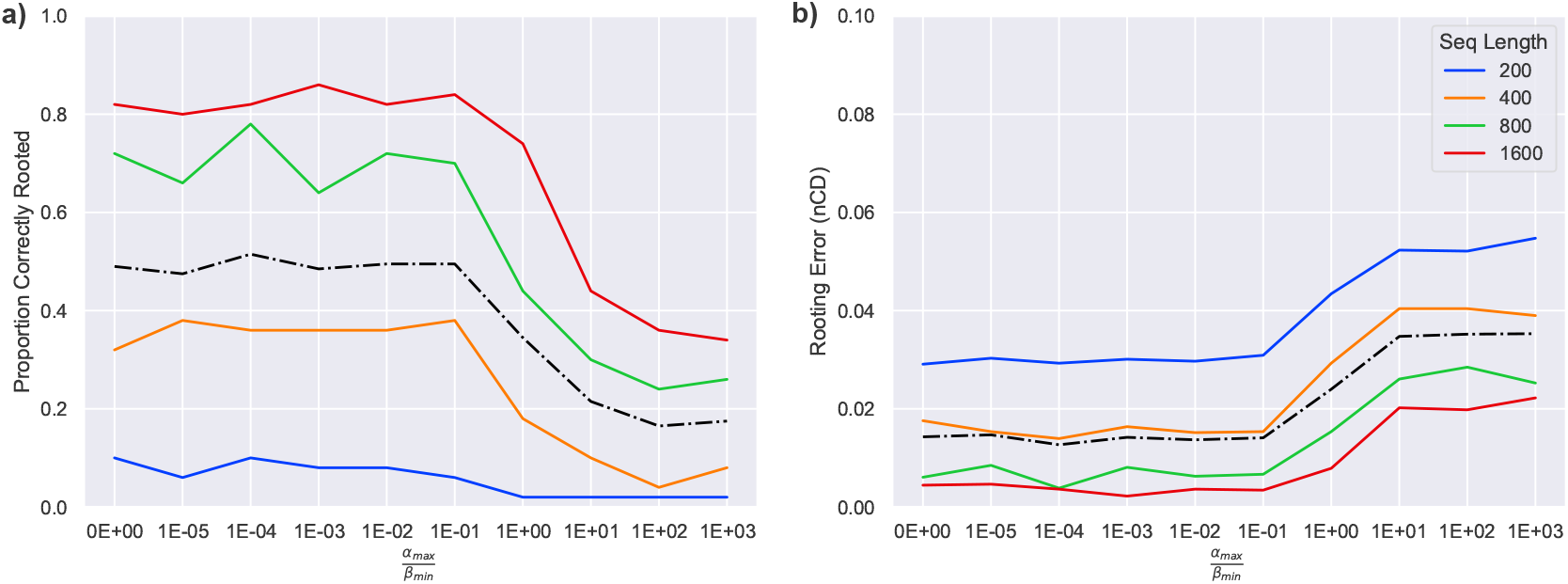
Impact of 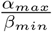 on QR-STAR. a) Proportion of the trees correctly rooted and b) rooting error (nCD) are shown for the S100 dataset from [35] with 50 replicates. The number of taxa is 101, the number of genes is 1000 and the average AD level is 0.46. The sequence length used to produce estimated gene trees varies between 200bp to 1600bp. The black dashed line corresponds to the average among sequence lengths. The value of 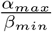 varies between 10^−5^ to 10^3^ in addition to 0. All *α* and *β* values are set as equal for all invariant or inequality penalty terms.

#### E.2 Additional Results for Comparing QR to QR-STAR

Figure E3 shows the result of rooting the model species tree with true gene trees on the S200 datasets. The trends are similar to Figure 4 in the main paper that uses estimated gene trees, except that in Figure 4, QR-STAR seems to be slightly less accurate than QR in the 500K, 1e-07 condition, although it’s more accurate in all model conditions of Figure E3. In general, the rooting error for all conditions (and especially the higher ILS conditions) seem to decrease when using true gene trees instead of estimated gene trees, which is expected.

**Fig. E3:**
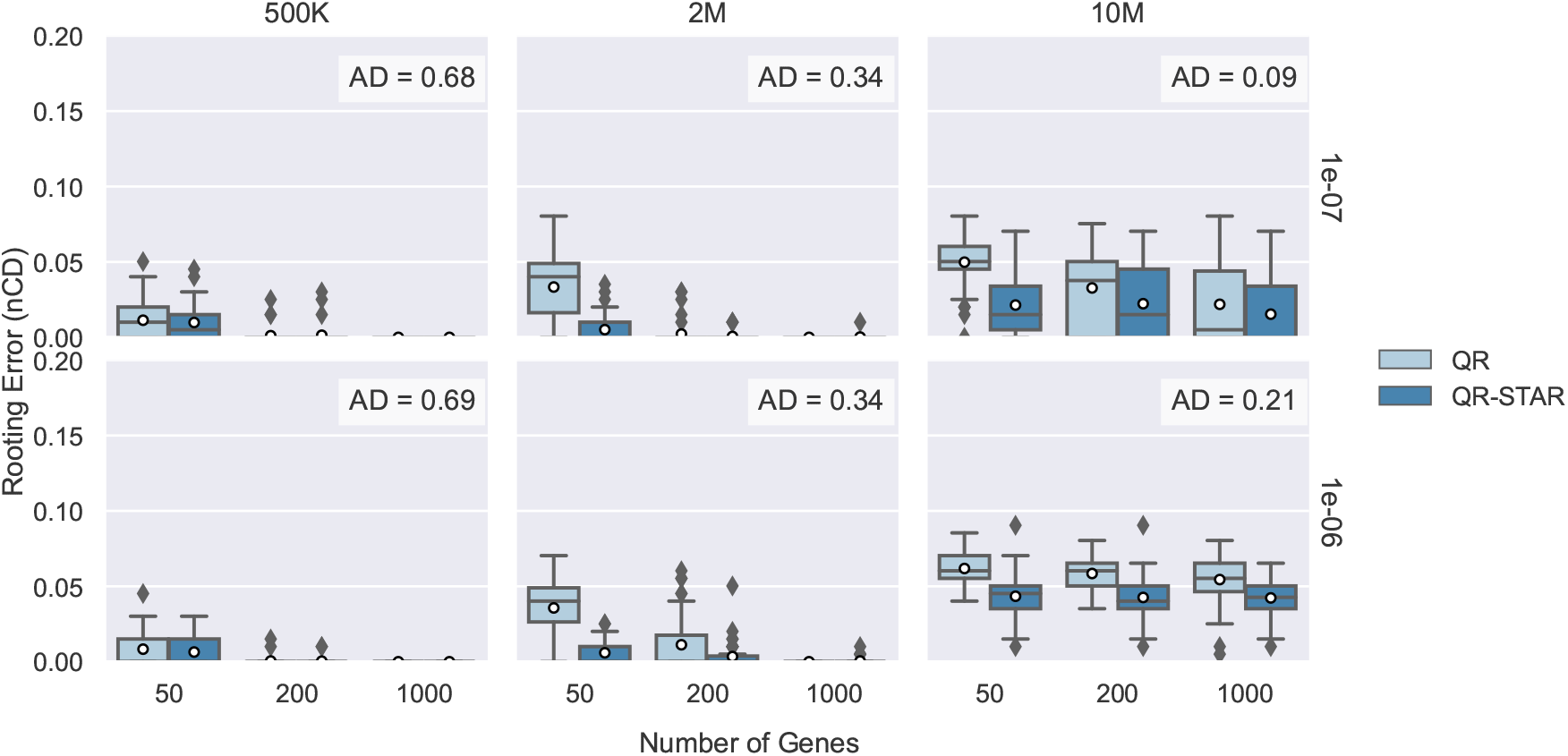
Rooting the model species tree with true gene trees on S200 datasets. Comparison between QR and QR-STAR in terms of rooting error (nCD) for rooting the true unrooted species tree topology using true gene trees on the 201-taxon datasets across 50 replicates. The columns show tree height (500K or high ILS, 2M or moderate ILS, and 10M or low ILS) and the rows show speciation rate (1e-06 for recent speciation, 1e-07 for deep speciation). QR-STAR is run with *C* =1e-02 and 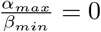.

Figures E4 and E5 show the overall error (measured in nCD) of an estimated rooted species tree which is the result of first estimating the unrooted species tree using ASTRAL and then rooting it using QR or QR-STAR. The difference that can be observed in these figures is that the error is generally higher when ILS rate is higher, especially for small number of genes, which is the opposite of the trend seen in Figures 4 and E3. However, it is well-known that ASTRAL degrades in accuracy as the ILS rates increase [35,19,20], and these figures suggest that the overall error in the rooted species tree is dominated by the species tree estimation error in the higher ILS conditions (refer to Sec. 6 for a discussion on this). The trends between QR and QR-STAR are similar to the previous figures, with QR-STAR always matching or improving upon QR, and the biggest improvement can be seen in the low-ILS settings.

**Fig. E4:**
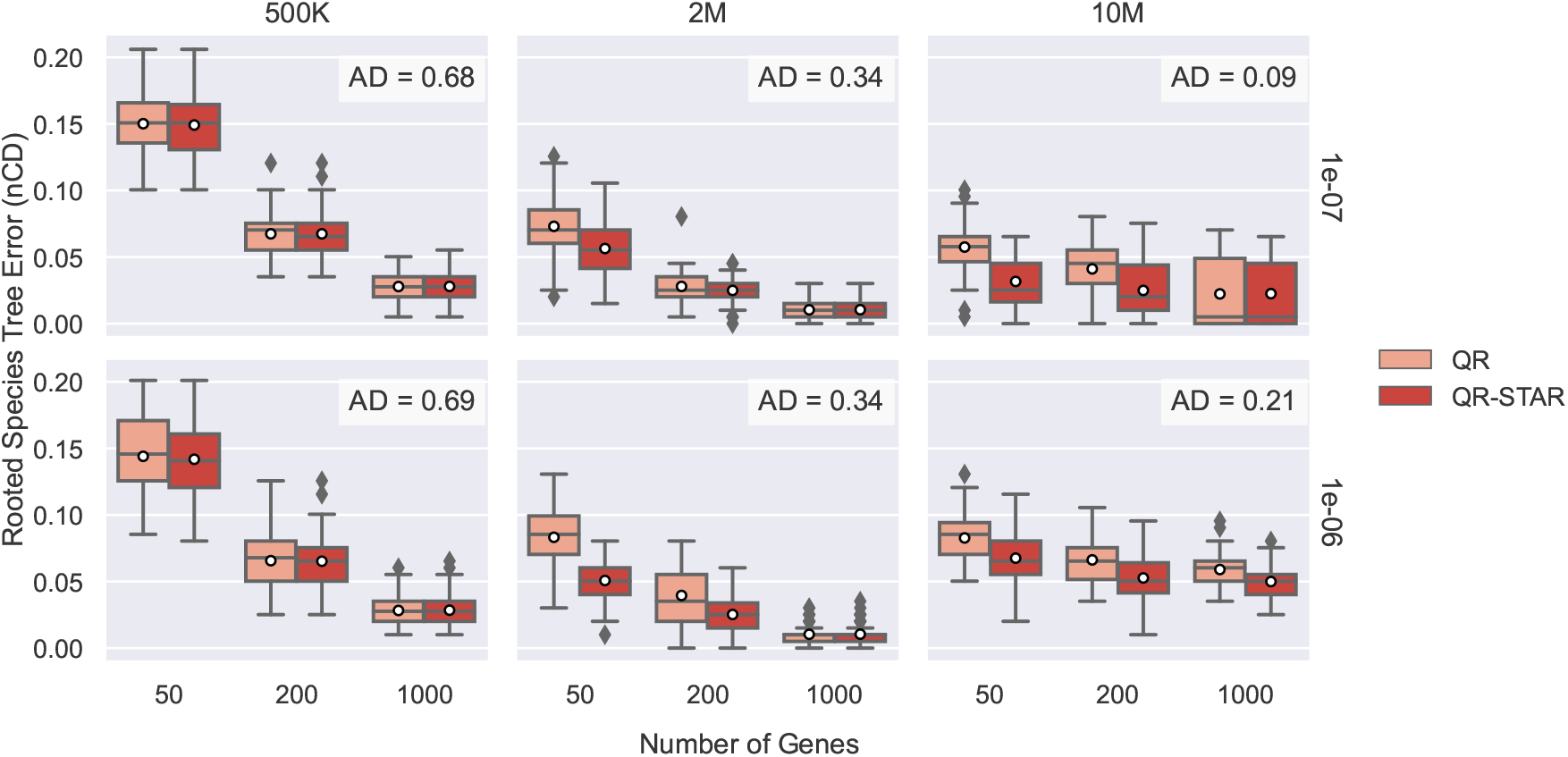
Rooting the ASTRAL species tree with true gene trees on S200 datasets. Comparison between QR and QR-STAR in terms of rooted species tree error (nCD) for rooting the species trees estimated by ASTRAL using true gene trees on the 201-taxon datasets across 50 replicates. The columns show tree height (500K or high ILS, 2M or moderate ILS, and 10M or low ILS) and the rows show speciation rate (1e-06 for recent speciation, 1e-07 for deep speciation). QR-STAR is run with *C* =1e-02 and 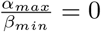.

**Fig. E5:**
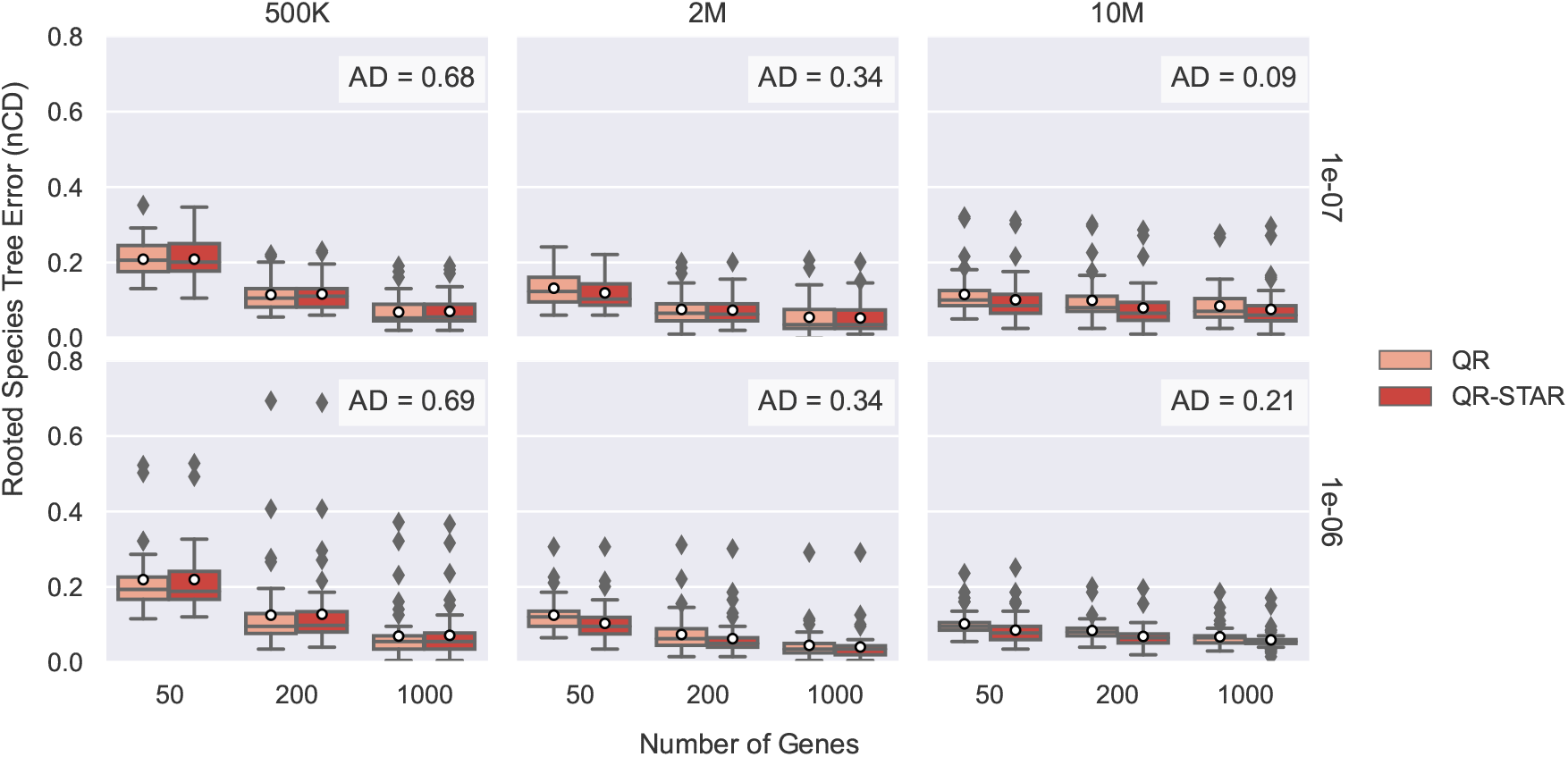
Rooting the ASTRAL species tree with estimated gene trees on S200 datasets. Comparison between QR and QR-STAR in terms of rooted species tree error (nCD) for rooting the species trees estimated by ASTRAL using estimated gene trees on the 201-taxon datasets across 50 replicates. The columns show tree height (500K or high ILS, 2M or moderate ILS, and 10M or low ILS) and the rows show speciation rate (1e-06 for recent speciation, 1e-07 for deep speciation). QR-STAR is run with *C* =1e-02 and 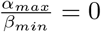.

### F Conflicts between 5-taxon Rooted Trees

**Fig. F6:**
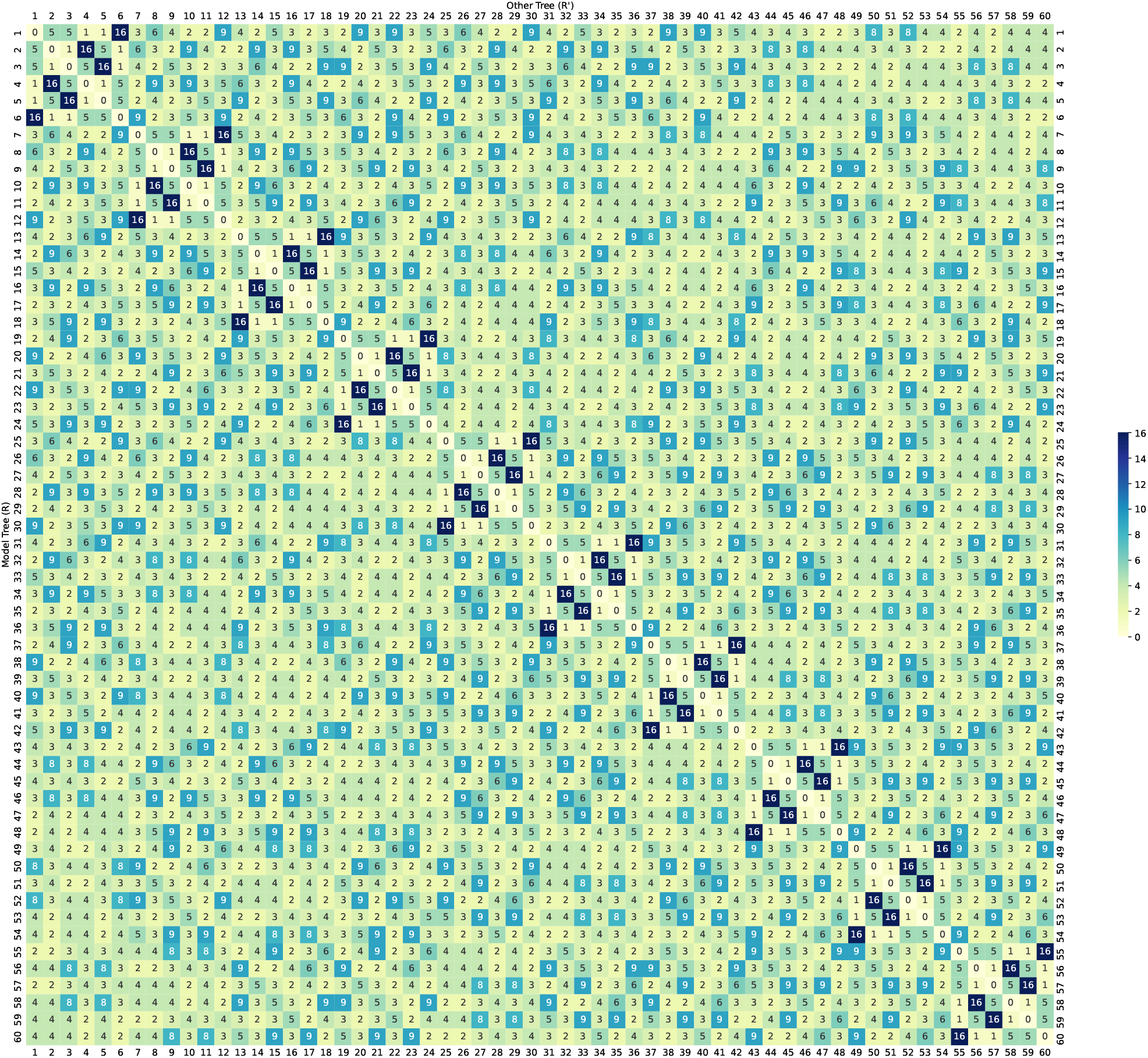
Conflicts between 5-taxon caterpillar trees. Heatmap showing the number of conflicting inequality penalty terms (the function |*V* (*R, R*^*′*^)|) for pairs of caterpillar 5-taxon rooted trees.

**Fig. F7:**
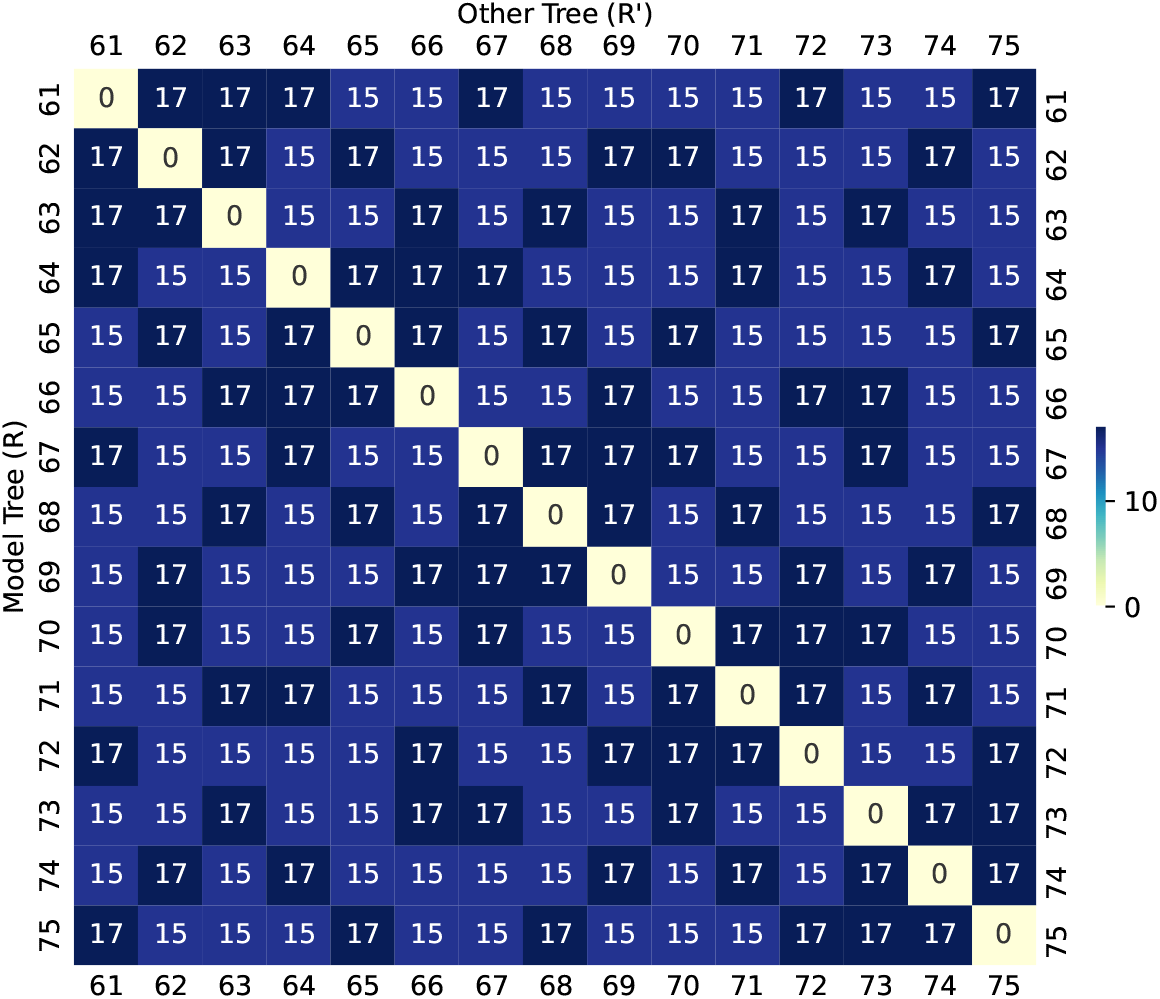
Conflicts between 5-taxon pseudo-caterpillar trees. Heatmap showing the number of conflicting inequality penalty terms (the function |*V* (*R, R*^*′*^)|) for pairs of pseudo-caterpillar 5-taxon rooted trees.

**Fig. F8:**
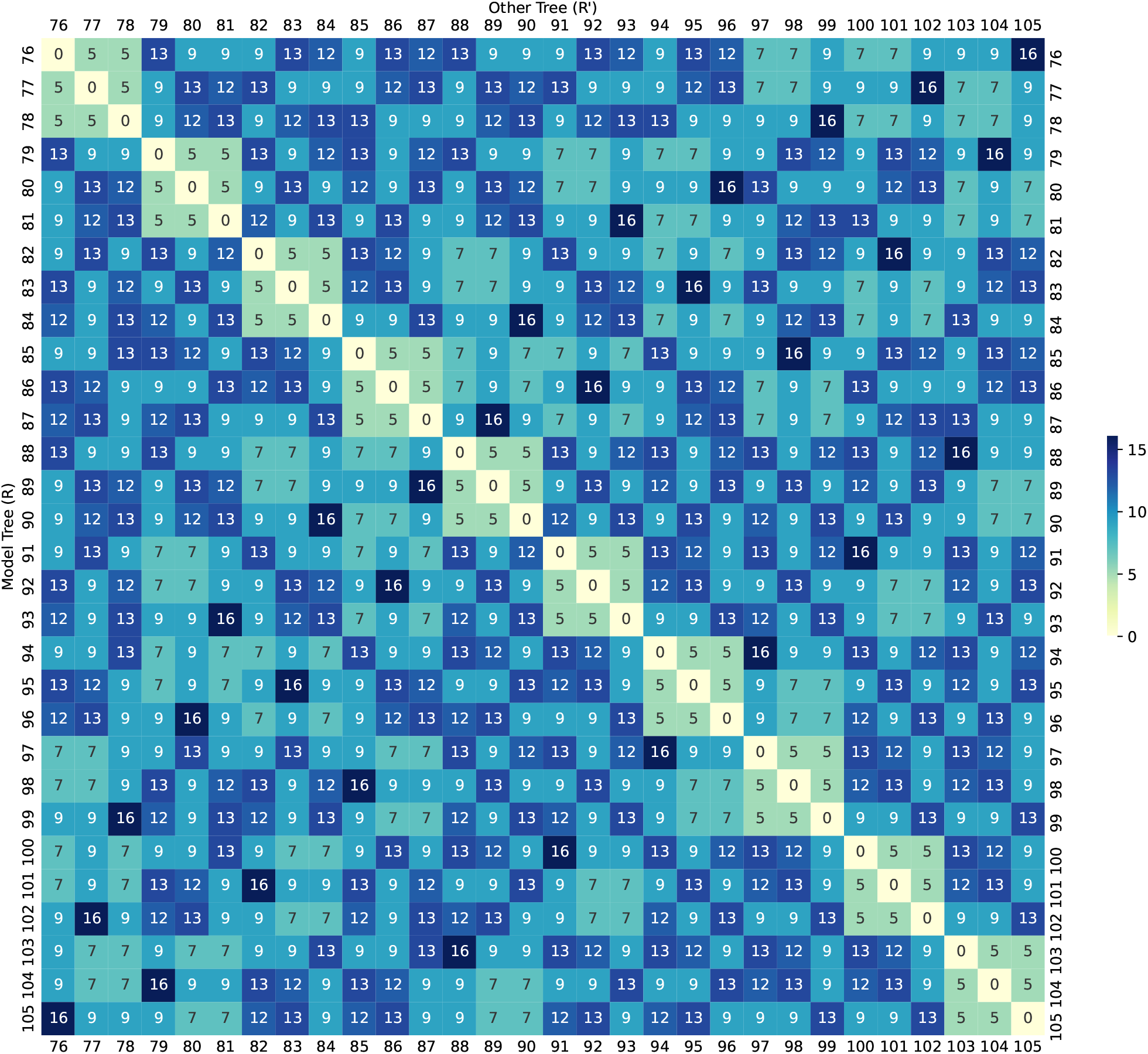
Conflicts between 5-taxon balanced trees. Heatmap showing the number of conflicting inequality penalty terms (the function |*V* (*R, R*^*′*^)|) for pairs of balanced 5-taxon rooted trees.

**Fig. F9:**
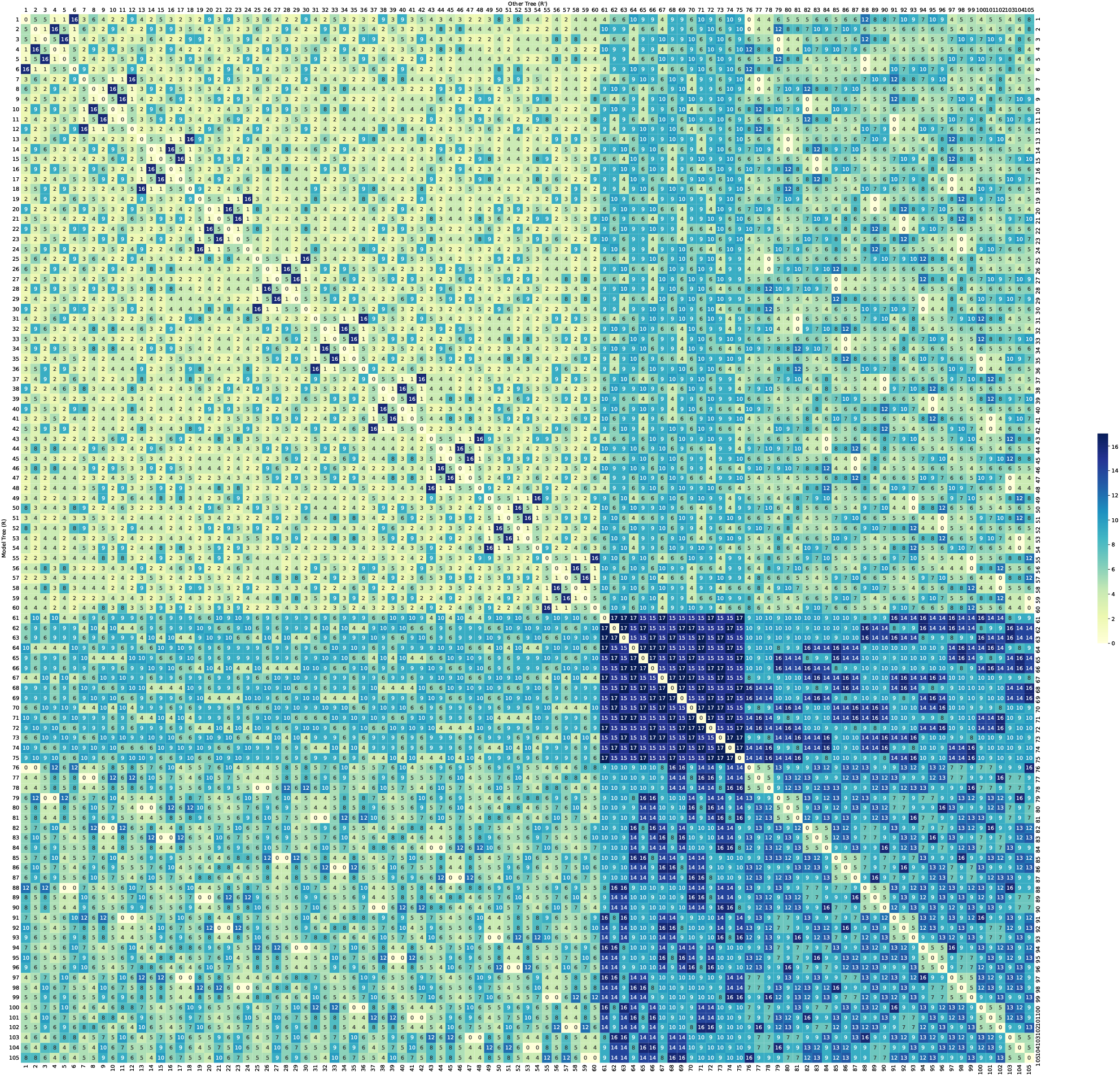
Conflicts between all 5-taxon rooted trees. Heatmap showing the number of conflicting inequality penalty terms (the function |*V* (*R, R*^*′*^)|) for all pairs of binary 5-taxon rooted trees.

## Footnotes

3 The labeling of rooted and unrooted trees in this paper is consistent with the notations and leaf-labeling used in Tables 4-5 in [2] as well as in [31].

4 Refer to Remark 1 for an explanation on why *α*_*a,b*_ does not need to be strictly positive.

## References

1. Alanzi, A.R., Degnan, J.H.: Inferring rooted species trees from unrooted gene trees using approximate Bayesian computation. Molecular Phylogenetics and Evolution 116, 13–24 (2017)

2. Allman, E.S., Degnan, J.H., Rhodes, J.A.: Identifying the rooted species tree from the distribution of unrooted gene trees under the coalescent. Journal of Mathematical Biology 62(6), 833–862 (2011)

3. Chan, Y.b., Li, Q., Scornavacca, C.: The large-sample asymptotic behaviour of quartet-based summary methods for species tree inference. Journal of Mathematical Biology 85(3), 1–22 (2022)

4. Emms, D.M., Kelly, S.: STRIDE: species tree root inference from gene duplication events. Molecular Biology and Evolution 34(12), 3267–3278 (2017)

5. Felsenstein, J.: Cases in which parsimony or compatibility methods will be positively misleading. Systematic Zoology 27(4), 401–410 (1978)

6. Graham, S.W., Olmstead, R.G., Barrett, S.C.: Rooting phylogenetic trees with distant outgroups: a case study from the commelinoid monocots. Molecular Biology and Evolution 19(10), 1769–1781 (2002)

7. Skarp-de Haan, C., Culebro, A., Schott, T., Revez, J., Schweda, E.K., Hänninen, M.L., Rossi, M.: Comparative genomics of unintrogressed Campylobacter coli clades 2 and 3. BMC Genomics 15(1), 1–14 (2014)

8. Hess, P.N., De Moraes Russo, C.A.: An empirical test of the midpoint rooting method. Biological Journal of the Linnean Society 92(4), 669–674 (2007)

9. Holland, B., Penny, D., Hendy, M.: Outgroup misplacement and phylogenetic inaccuracy under a molecular clock—a simulation study. Systematic Biology 52(2), 229–238 (2003)

10. Hudson, R.R.: Testing the constant-rate neutral allele model with protein sequence data. Evolution pp. 203–217 (1983)

11. Jun, S.R., Leuze, M.R., Nookaew, I., Uberbacher, E.C., Land, M., Zhang, Q., Wanchai, V., Chai, J., Nielsen, M., Trolle, T., Lund, O., Buzard, G.S., Pedersen, T.D., Wassenaar, T.M., Ussery, D.W.: Ebolavirus comparative genomics. FEMS Microbiology Reviews 39(5), 764–778 (2015)

12. Larget, B.R., Kotha, S.K., Dewey, C.N., Ané, C.: BUCKy: gene tree/species tree reconciliation with Bayesian concordance analysis. Bioinformatics 26(22), 2910–2911 (2010)

13. Li, C., Matthes-Rosana, K.A., Garcia, M., Naylor, G.J.: Phylogenetics of Chondrichthyes and the problem of rooting phylogenies with distant outgroups. Molecular Phylogenetics and Evolution 63(2), 365–373 (2012)

14. Liu, L., Yu, L., Edwards, S.V.: A maximum pseudo-likelihood approach for estimating species trees under the coalescent model. BMC Evolutionary Biology 10(1), 1–18 (2010)

15. Maddison, W.P.: Gene trees in species trees. Systematic Biology 46(3), 523–536 (1997)

16. Maddison, W.P., Donoghue, M.J., Maddison, D.R.: Outgroup analysis and parsimony. Systematic Biology 33(1), 83–103 (1984)

17. Mahbub, M., Wahab, Z., Reaz, R., Rahman, M.S., Bayzid, M.S.: wQFM: highly accurate genome-scale species tree estimation from weighted quartets. Bioinformatics 37(21), 3734–3743 (2021)

18. Mai, U., Sayyari, E., Mirarab, S.: Minimum variance rooting of phylogenetic trees and implications for species tree reconstruction. PloS One 12(8), e0182238 (2017)

19. Mirarab, S., Reaz, R., Bayzid, M.S., Zimmermann, T., Swenson, M.S., Warnow, T.: ASTRAL: genome-scale coalescent-based species tree estimation. Bioinformatics 30(17), i541–i548 (2014)

20. Mirarab, S., Warnow, T.: ASTRAL-II: coalescent-based species tree estimation with many hundreds of taxa and thousands of genes. Bioinformatics 31(12), i44–i52 (2015)

21. Mitzenmacher, M., Upfal, E.: Probability and computing: Randomization and probabilistic techniques in algo-rithms and data analysis. Cambridge University Press (2017)

22. Molloy, E.K., Warnow, T.: To include or not to include: the impact of gene filtering on species tree estimation methods. Systematic Biology 67(2), 285–303 (2018)

23. Posada, D.: Phylogenomics for systematic biology. Systematic Biology 65(3), 353–356 (2016)

24. Price, M.N., Dehal, P.S., Arkin, A.P.: FastTree 2–approximately maximum-likelihood trees for large alignments. PloS One 5(3), e9490 (2010)

25. Rabiee, M., Mirarab, S.: QuCo: quartet-based co-estimation of species trees and gene trees. Bioinformatics 38(Supplement 1), i413–i421 (2022)

26. Renner, S.S., Grimm, G.W., Schneeweiss, G.M., Stuessy, T.F., Ricklefs, R.E.: Rooting and dating maples (Acer) with an uncorrelated-rates molecular clock: implications for North American/Asian disjunctions. Systematic Biology 57(5), 795–808 (2008)

27. Robinson, D.F., Foulds, L.R.: Comparison of phylogenetic trees. Mathematical Biosciences 53(1-2), 131–147 (1981)

28. Rosenberg, N.A.: Counting coalescent histories. Journal of Computational Biology 14(3), 360–377 (2007)

29. Shekhar, S., Roch, S., Mirarab, S.: Species tree estimation using ASTRAL: how many genes are enough? IEEE/ACM Transactions on Computational Biology and Bioinformatics 15(5), 1738–1747 (2017)

30. Simmons, M.P., Springer, M.S., Gatesy, J.: Gene-tree misrooting drives conflicts in phylogenomic coalescent analyses of palaeognath birds. Molecular Phylogenetics and Evolution 167, 107344 (2022)

31. Tabatabaee, Y., Sarker, K., Warnow, T.: Quintet Rooting: rooting species trees under the multi-species coalescent model. Bioinformatics 38(Supplement 1), i109–i117 (2022)

32. Tria, F.D.K., Landan, G., Dagan, T.: Phylogenetic rooting using minimal ancestor deviation. Nature Ecology & Evolution 1(1), 1–7 (2017)

33. Wascher, M., Kubatko, L.: Consistency of SVDQuartets and maximum likelihood for coalescent-based species tree estimation. Systematic Biology 70(1), 33–48 (2021)

34. Willson, J., Tabatabaee, Y., Liu, B., Warnow, T.: DISCO+QR: Rooting species trees in the presence of GDL and ILS. bioRxiv (2023), doi: https://doi.org/10.1101/2023.01.02.522492

35. Zhang, C., Rabiee, M., Sayyari, E., Mirarab, S.: ASTRAL-III: polynomial time species tree reconstruction from partially resolved gene trees. BMC Bioinformatics 19(6), 15–30 (2018)

